# Molecular analysis of network vulnerability to α-synuclein pathology reveals PAKs as therapeutic targets for Parkinson’s disease

**DOI:** 10.1101/2024.10.22.619411

**Authors:** Naman Vatsa, Julia K. Brynildsen, Thomas M. Goralski, Kevin Kurgat, Lindsay Meyerdirk, Libby Breton, Daniella DeWeerd, Laura Brasseur, Lisa Turner, Katelyn Becker, Kristin L. Gallik, Christine Isaguirre, Ryan D. Sheldon, Dani S. Bassett, Michael X. Henderson

## Abstract

α-Synuclein misfolding and progressive accumulation drive a pathogenic process in Parkinson’s disease, yet many brain regions develop more or less pathology than would be predicted by connectivity alone, indicating that intrinsic biological factors influence regional vulnerability. Here, we combined whole brain mapping of α-synuclein pathology in wildtype mice 3 days to 9 months after seeding with network diffusion modeling based on anatomical connectivity to derive quantitative measures of regional vulnerability. To identify molecular drivers of this vulnerability, we generated a brain-wide sPAtial Neuron Gene Expression Atlas (PANGEA) and compared regional transcriptional profiles with model-derived vulnerability scores. Vulnerable regions were enriched for specific cellular programs. Specific kinases were also enriched in vulnerable regions, leading to the identification of group II p21-activated kinases (PAKs) as candidate regulators of α-synucleinopathy. Pharmacological inhibition of group II PAKs reduced α-synuclein aggregation and protected against neuron loss in primary neurons, remained effective when administered after pathology initiation, and suppressed pathology *in vivo.* Consistent with these findings, genetic depletion of PAK5/6 also suppressed α-synuclein aggregation. Together, these results establish a framework linking network-level measures to cellular mechanisms of selective vulnerability and nominate group II PAKs as promising disease-modifying therapeutic targets for Parkinson’s disease.

## INTRODUCTION

Parkinson’s disease (PD) is a progressive disorder characterized by loss of dopaminergic neurons and accumulation of α-synuclein Lewy pathology throughout the brain^1,2^. Extensive cellular dysfunction has been documented in PD as well as in cell and animal models, with prominent involvement of mitochondria, lysosomes, and synapses^3^. However, PD manifests as a network disorder^4^. While dopaminergic neurons are lost in the substantia nigra pars compacta, movement symptoms are manifested due to disruption of basal ganglia control of motor cortical output. People with PD also exhibit a range of non-motor symptoms which are attributed to dysfunction in many other central and peripheral nervous system circuits^5^. α-Synuclein Lewy pathology is present in these disrupted circuits, and recent evidence suggests that misfolded α-synuclein may itself progress through synaptic connections, acting as the substrate of progressive dysfunction^5,6^.

Symptomatic therapies for PD, such as dopamine replacement and deep brain stimulation, have sought to restore function in disrupted neural networks^4^. In contrast, disease-modifying therapies have largely been identified based on cellular mechanisms^7^, with limited understanding of how these interventions influence network-level dysfunction. This disconnect arises in part because genes disrupted in disease can be readily linked to their encoded proteins, enabling high-throughput investigation of cellular dysfunction. However, translating these findings to the level of large-scale neural circuitry remains challenging. Another barrier to identifying network-level targets is the challenge of characterizing pathological network disruptions that occur in PD, as these can typically only be assessed at disease end-stage.

Neuroimaging has arisen as one method to assay longitudinal measurements of pathological outcomes. In addition to structural magnetic resonance imaging (MRI) and neurotransmitter receptor imaging, one can directly measure the regional residence of pathological proteins with positron emission tomography (PET) tracers. This technology currently exists for amyloid β and tau pathologies^8^, and a variety of mathematical models have been used to predict pathology patterns using anatomical connectivity, functional connectivity, or regional proximity to form the base network^9–23^.

While PET tracers are currently still being optimized for α-synuclein inclusions^24–26^, we and others have leveraged network diffusion modeling to understand pathology progression in seed-based animal models. In these models, small amounts of misfolded α-synuclein or tau are injected into the brains of wildtype mice^27,28^. The subsequent propagation of pathology is constrained by anatomical connectivity^29–38^. However, connectivity alone is not sufficient to predict pathology in all regions. Cell autonomous vulnerability likely underlies differences in model predictivity, and we can use the residual variance in a connectivity-based model to estimate regional vulnerability^39^.

In the current study, we employed a network-based approach to identify novel targets for PD, followed by cellular and *in vivo* validation to confirm their potential. While we have previously established the ability to generate regional vulnerability measures using network modeling^29^, efforts to translate this into molecular insight were impinged by two main factors—low pathology resolution (134 brain regions) and limited regional gene expression data (4277 genes). We have now implemented a user-guided workflow to generate quantitative pathology measures in 1046 brain regions at 8 timepoints in mice injected with α-synuclein pre-formed fibrils. Regional pathology burden was well-described by linear diffusion through anatomical connections, and residuals in this model were used to develop a measure of relative regional vulnerability. We also developed a sPAtial Neuronal Gene Expression Atlas (PANGEA) with coverage of 19,781 genes in 302 of the 722 gray matter regions in the Allen Brain Atlas. We compared regional vulnerability to PANGEA to identify gene expression patterns related to vulnerability. Interestingly, we identified gene sets associated with both resilience (oxidative phosphorylation) and vulnerability (complement/coagulation cascade).

By cross-referencing kinases associated with vulnerability to those whose expression is altered in neurons with Lewy pathology^40^, we identified 12 kinases with a possible association with cellular vulnerability. We then screened those kinases and identified p21-activated kinases (PAKs) as modifiers of α-synucleinopathy in primary neurons. We isolated this effect to group II PAKs and showed that group II PAK inhibition reduces α-synuclein pathology and rescues neuron death. Genetic depletion of PAK5 and PAK6 also reduced α-synuclein pathology, further supporting a role for group II PAKs in α-synucleinopathy. Finally, we demonstrated that group II PAK inhibition *in vivo* was sufficient to reduce α-synuclein pathology in a mouse model.

Together, these results demonstrate the power of network-level analysis to reveal molecular insight and identify group II PAKs as potential targets for therapeutic development in PD.

## RESULTS

### α-Synuclein PFFs induce a broad pathology pattern, influenced by regional neuron types

We have previously hypothesized that anatomical patterns of protein pathology arise from a combination of intrinsic cellular vulnerability and network connectivity^39^. To test this hypothesis, we needed to develop high-resolution maps of α-synuclein pathology progression over time, computational prediction of pathology based on anatomical connectivity, a brain-wide regional gene expression map, and cellular validation of network-derived vulnerability measures. To induce α-synuclein pathology, we injected 3-month-old wildtype mice with α-synuclein pre-formed fibrils (PFFs) in the dorsal striatum (**Fig. 1A**). Mice were aged 0.1, 0.2, 0.3, 0.5, 1, 3, 6, or 9 months post-injection (MPI) to capture the progression of pathology. Each brain was sampled systematically through coronal sectioning, registered to the Allen Brain Atlas (ABA) CCFv3 and segmented for pathology.

**Figure 1.**
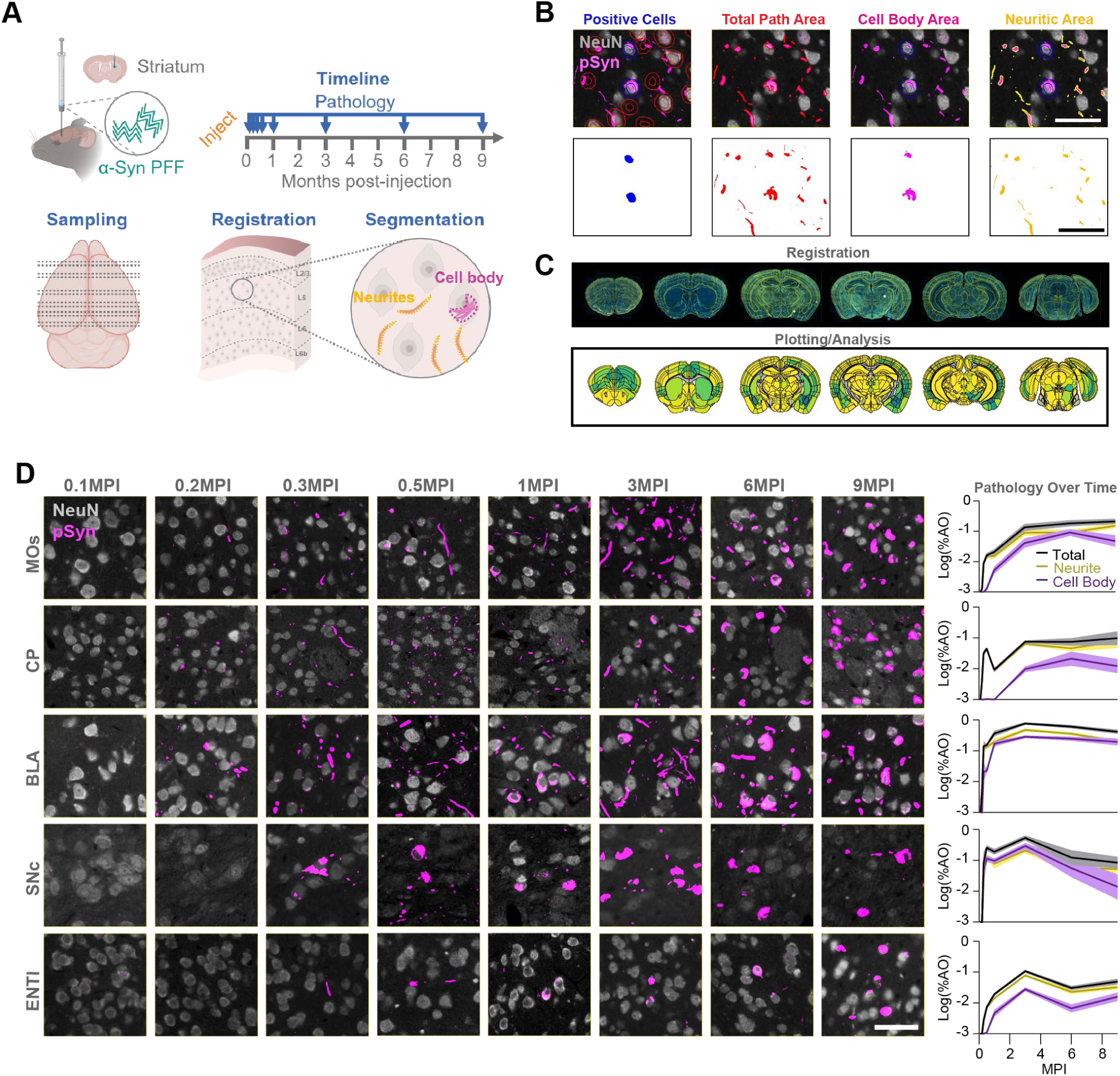
α-Synuclein PFFs induce a broad pathology pattern, influenced by regional neuron types. (A) Experimental schematic. Wildtype mice were injected in the dorsal striatum with α-synuclein pre-formed fibrils (PFFs). Mice were aged a further 0.1, 0.2, 0.3, 0.5, 1, 3, 6, or 9 months post-injection (MPI). Coronal sections representing the majority of regions with pathology were routinely sectioned for all mice. Sections were stained and registered to the Allen Brain Atlas (ABA) CCFv3 and multiple forms of pathology were segmented and quantified. (B) Example of pathology segmentation from mouse brain tissue stained for neurons (NeuN) and α-synuclein pathology (pSyn). The top panels show how the segmentation performs on the tissue while the lower panels are the resulting pathology masks. Scale bar = 50 μm. (C) Example set of sections registered to the ABA (top panel) and the resulting quantitative pathology plots (bottom panel). (D) Representative α-synuclein pathology staining across timepoints for 5 different brain regions. Plots on the right represent the total, cell body, or neuritic pathology quantification for those regions. Scale bar = 50 μm.

α-Synuclein PFF injection induces broad neuritic and cell body inclusions positive for pS129 α-synuclein, similar to PD brains. While cell body inclusions represent the impact of pathology in a particular region, neuritic pathology could also be present in neurons projecting to or through a region. Therefore, following staining of tissue for pS129 α-synuclein and NeuN for neuronal cell bodies, we developed segmentation strategies to capture the number of neurons with cell body inclusions or the percentage area occupied of total, cell body, or neuritic pathology (**Fig. 1B, S1A**). Each section was registered to the ABA CCFv3 using a suite of software known as the QUINT workflow^41^ (**Fig. S1B**). We adapted this workflow to include our segmentation measures^42^ and developed an accompanying plotting and statistical analysis platform for registered datasets (**Fig. 1C**).

We demonstrate representative pathology progression in several regions that captures some of the major patterns observed across the brain (**Fig. 1D, S2**). Pathology develops rapidly within motor cortex and amygdala as early as 0.2 MPI, with neuritic pathology preceding cell body pathology. Despite direct proximity to the injection site, the caudoputamen is relatively slow to develop pathology. Even when pathology does develop there, it is typically neuritic until later timepoints, suggesting that the GABAergic neurons in the caudoputamen are resistant to developing α-synuclein pathology. In contrast, the substantia nigra pars compacta rapidly develops cell body inclusions, and these inclusions are lost at later timepoints, consistent with death of those neurons^31^.

### Time- and region-dependent progression of α-synuclein pathology

We were able to map total, cell body, and neuritic α-synuclein pathology in 1046 regions of the brain at the 8 timepoints (**Fig. 2A**, **S3**, **S4A**-**S4C**). The pathology patterns observed in individual regions are also evident at the whole-brain level. Total pathology peaks by 3 MPI and plateaus thereafter (**Fig. 2B**). Some of that leveling off likely relates to consolidation of pathology into more compact somatic inclusions. Consistent with this hypothesis, cell body pathology is delayed compared to neuritic pathology, but continues to climb at later timepoints (**Fig. 2B**). Globally, pathology is initially more lateralized, with ipsilateral regions bearing more pathology. Later, pathology becomes more broadly distributed bilaterally (**Fig. 2C**). Finally, we examined the timepoints at which pathology peaks in different regions (**Fig. 2D**, **S4D**, **S4E**). While there is a global peak of pathology at 3 MPI, this is largely driven by amygdala, midbrain and caudal cortical regions. Frontal regions mostly peak at 6 or 9 MPI.

**Figure 2.**
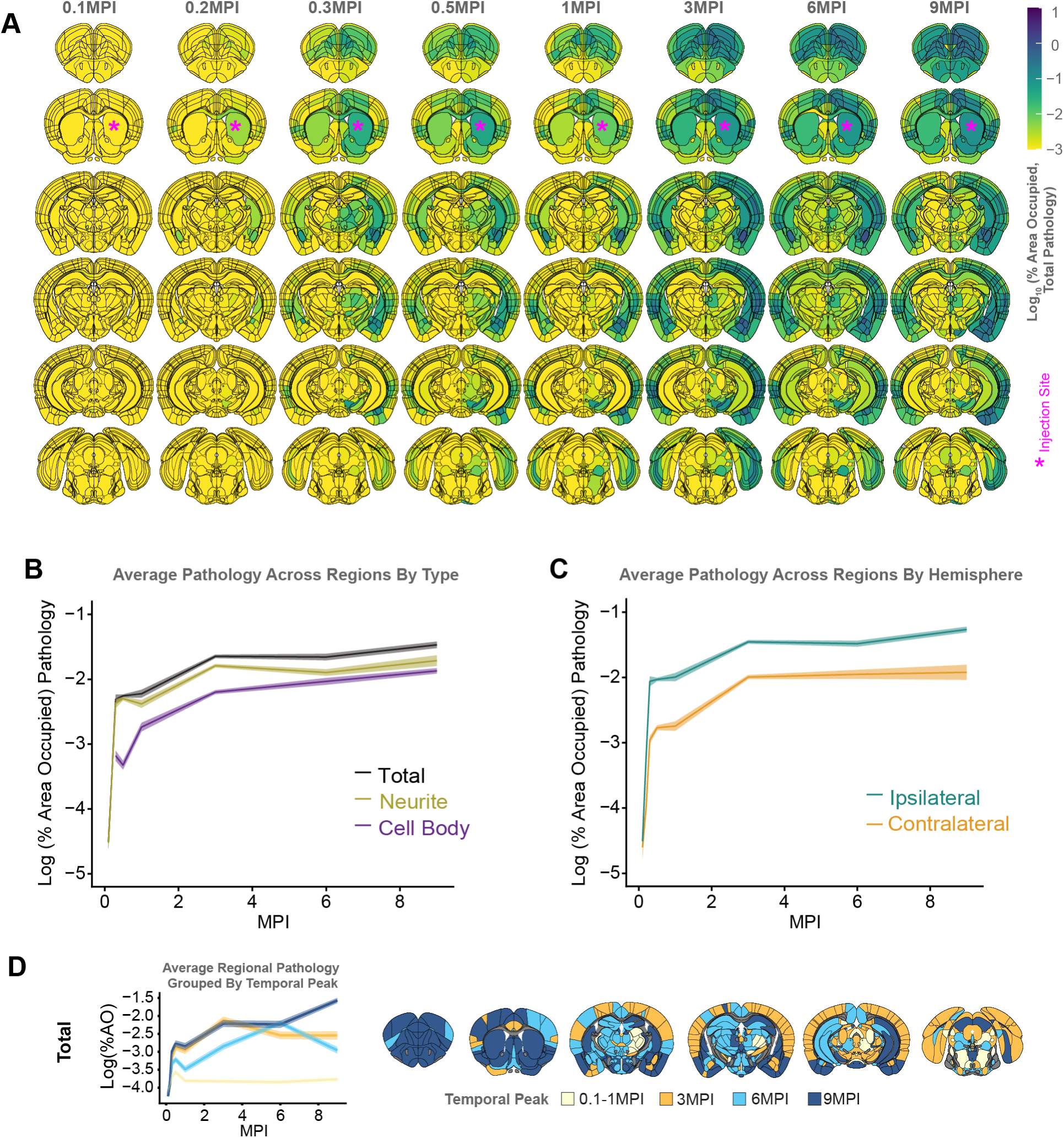
Time- and region-dependent progression of α-synuclein pathology. (A) Pathology heatmaps representing the average pathology in each anatomical brain region at the corresponding timepoint. *Injection sites. (B) Plot of neurite, cell body, and total pathology, summed across all regions in both hemispheres and averaged across mice at each time point, demonstrates the predominance of neuritic pathology, especially early, and the overall plateau of pathology at 3 MPI. (C) Plot of total pathology, summed across regions within each hemisphere and averaged across mice at each time point, demonstrates the early predominance of ipsilateral pathology and later gains in contralateral pathology. (D) Total α-synuclein pathology peaks at different times in specific regional groups. The average pathology level of regions that peak at different times is plotted as a group in the panel on the left. For visualization purposes, regions containing 0 pathology were represented as –4 on the log scale.

**Figure 3.**
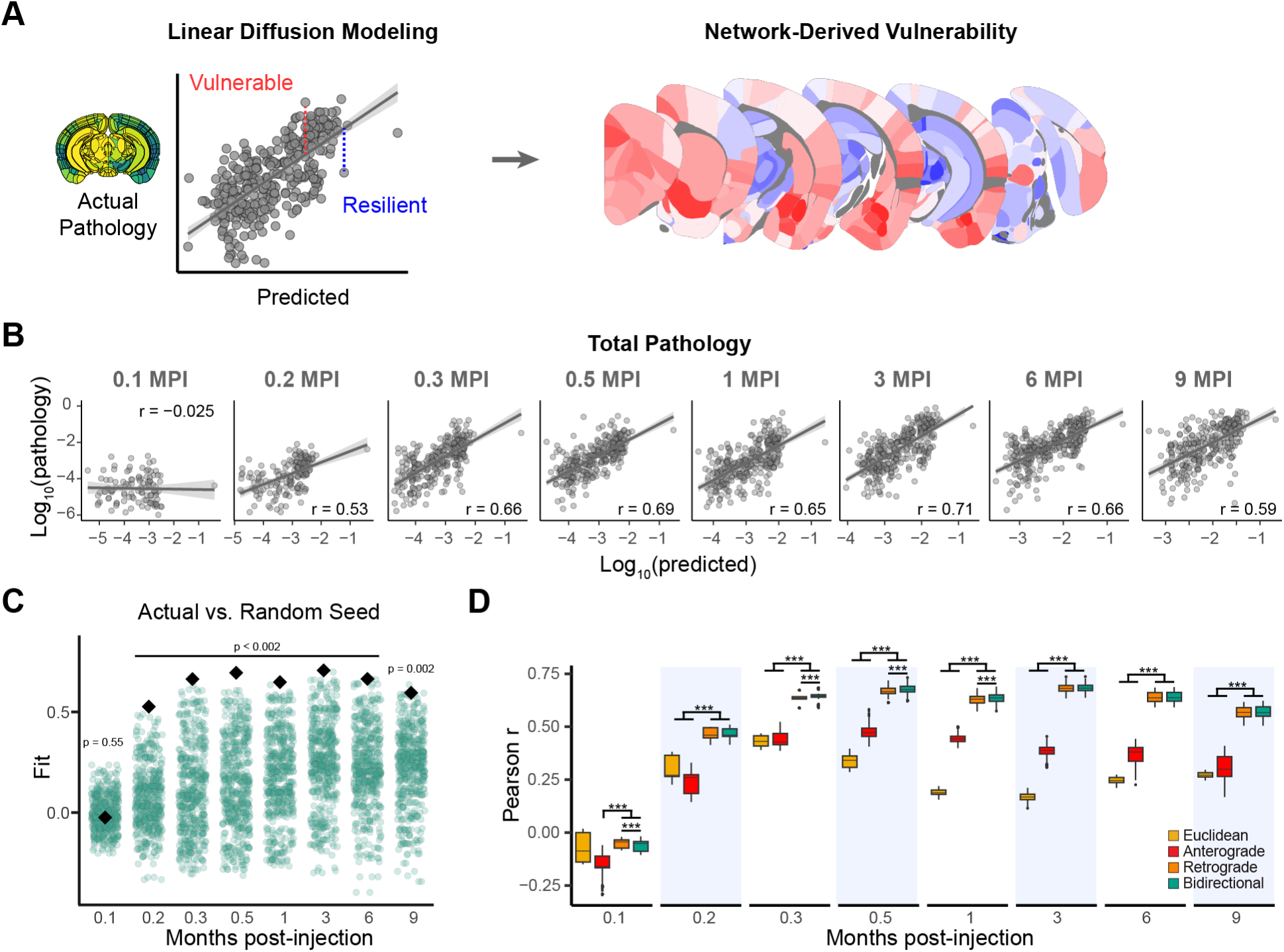
Computational model of pathology based on anatomical connectivity shows predictivity of regional α-synuclein pathology. (A) Schematic illustrating how regional vulnerability to pathology is derived from predictions generated by the linear network diffusion model. (B) Predictions of total α-synuclein pathology from linear network diffusion models based on bidirectional (anterograde and retrograde) anatomical connections. Solid lines represent the line of best fit, and shading represents 95% confidence intervals. Each dot represents a different brain region where α-synuclein pathology was measured. (C) Comparison of Pearson’s *r* values obtained by fitting bidirectional spread models using actual (black diamond) and alternate (green circles) seed regions. (D) Comparison of Euclidean, anterograde, retrograde, and bidirectional model fits across 500 held-out samples derived from 500 iterations of a cross-validation process. Retrograde and bidirectional spread models performed significantly better than the anterograde spread model at all time points, and significantly better than the Euclidean spread model at all time points except 0.1 MPI (\*\*\**P* < 0.002).

To assess whether our coronal (2D) sampling strategy accurately captured brain-wide pathology patterns, one brain was fixed at 3 MPI, optically cleared, and stained for pS129 α-synuclein. The whole brain was imaged (3D) and registered to the ABA CCFv3. Segmentation of total pathology volume was performed as a direct comparison to pathology area captured in coronal sections (**Fig. S5A**, **S5B**). Of note, imaging quality was insufficient with this modality to confidently distinguish between cell body and neuritic inclusions, so only total inclusion area was measured. Overall, there was a strong correlation between 2D and 3D datasets (**Fig. S5C-S5D**). However, there were some discrepancies between the two datasets that corresponded to major anatomical divisions. It was apparent that some of the over-representation in the 3D dataset was related to non-specific staining of ventricles, including those in the hypothalamus, and misregistration of regions (**Fig. S5E, S5F**). For example, the substantia nigra is a very thin nucleus in the ABA CCFv3 and the 3D registration misaligned the substantia nigra pars compacta, assigning the pathology in this region to other midbrain nuclei. Cortical laminar alignment was also sub-optimal in the 3D registration (**Fig. 4G, 4H**). We did identify some thalamic regions in the 3D dataset that were not well-sampled in the 2D dataset, partially because those nuclei are broadly distributed along the rostro-caudal axis and may not have been enriched in the assessed coronal samples (**Fig. S5I, S5J**). Overall, the comparison of 2D and 3D datasets highlighted the brain-wide coverage achieved with coronal sampling and the greater opportunity for manual quality control during brain registration.

**Figure 4.**
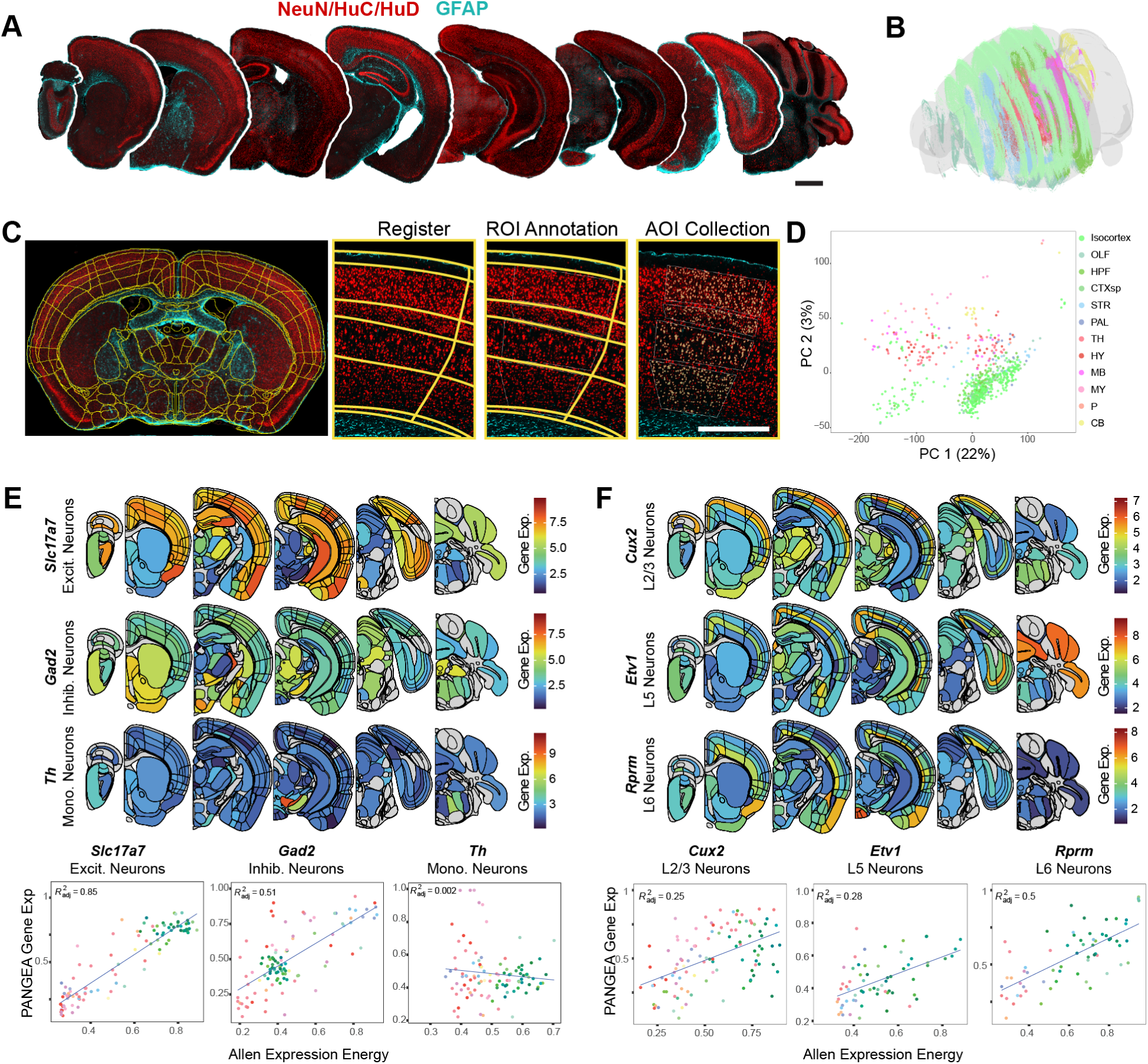
sPAtial Neuronal Gene Expression Atlas (PANGEA) (A) Representative coronal sections throughout the rostro-caudal axis of the mouse brain stained with NeuN/HuC/HuD (red) and GFAP (cyan). Scale bar = 1 mm. (B) Transparent 3D brain reconstruction of all sections sampled for this study placed in the appropriate 3D space. Made with MeshView. (C) Example segmented brain section on the GeoMx instrument. Subsequent panels demonstrate zoomed view of the registration, region-of-interest (ROI) annotation, and area-of-illumination (AOI) for the collection of regional neuronal transcripts. Scale bar = 0.5 mm. (D) Principal component analysis plot with each AOI plotted as a single dot colored by the major anatomical region. Regions show segregation mainly dependent on the major brain region. (E, F) Anatomical heatmaps and correlation plots comparing PANGEA data to Allen *in situ* hybridization expression energy data^47^. PANGEA gene expression related to neuron types shows the expected regional pattern from previous studies and correlates well with Allen expression energy, except for *Th*, which is very sparsely expressed. Correlation plot coloring is based on ABA anatomical region coloration (see legend on panel D).

Overall, the pathology in this model is progressive and relates to anatomical connectivity of the injection site, but with clear influence of regional vulnerability as well. To further assess these patterns and develop a measure of regional vulnerability, we turned to computational network modeling.

### Computational model of pathology based on anatomical connectivity shows predictivity of regional α-synuclein pathology

In previous studies, linear network diffusion models have been shown to explain the spread of pathological tau and α-synuclein proteins along the brain’s structural connectome^29–38^. These studies have determined that patterns of pathology occupancy are primarily driven by spread in the retrograde direction. Here, we build on this prior work by modeling the spread of pathology from high spatiotemporal resolution data, including both cell body and neuritic pathology. We hypothesized that modeling the extent of pathology explained by connectivity would enable us to derive variation not explained by connectivity, which we refer to broadly as regional vulnerability (**Fig. 3A**). Consistent with our previous analysis that includes hundreds of annotated regions, the current analysis of higher resolution data yielded a strong fit to actual pathology measures, as early as 0.2 MPI (**Fig. 3B**).

To evaluate model specificity to the injection site, we randomly selected 500 alternate seed regions and recomputed model fit using each of these regions. The experimental injection site resulted in a stronger correlation between actual and predicted pathology at every time point except the earliest (0.1 MPI), further supporting the specificity of the model fit to the connectivity from the striatum (**Fig. 3C**).

To validate our choice of the bidirectional network diffusion model, we statistically compared its out-of-sample performance to models based on retrograde spread alone, anterograde spread alone, and Euclidean distance. The retrograde and bidirectional spread models performed significantly better than the Euclidean distance model at all time points except the first (0.1 MPI) and outperformed the anterograde spread model at every time point (**Fig. 3D**). The bidirectional spread model outperformed the retrograde spread model only slightly at 0.3, 0.6, and 1 MPI. Although our model selection was informed by earlier reports demonstrating that spread predominantly occurs in the retrograde direction, this comparison provided a direct assessment of its explanatory strength relative to alternative models. Further, evaluating out-of-sample performance confirmed the generalizability of our model.

Consistent with the observation that neuritic pathology dominates at early time points, we observed strong correlations between actual and model-predicted neuritic pathology beginning at 0.2 MPI (**Fig. S6A**). In contrast, there was insufficient cell body pathology to fit a model until 0.3 MPI, and model fit remained weak until 3 MPI (**Fig. S6B**). This discrepancy likely arises from neuritic pathology developing rapidly while cell body pathology develops slowly, resulting in fewer regions with pathology to properly estimate model fit. In both cell bodies and neurites, the strongest fit was observed at 3 MPI. Comparison of the actual seed site with alternate seed sites revealed a similar temporal pattern, with strong discrimination of the actual seed site for neuritic pathology from 0.2 MPI onward (**Fig. S6C**), and for cell body pathology from 3 MPI onward (**Fig. S6D**).

Although the linear network diffusion model parameters were fit to a caudoputamen injection site, we explored the utility of this model in determining α-synuclein pathology progression following injection into other sites. Model parameters remained consistent with those used for the caudoputamen injection site, but the initiation site was shifted throughout the rostro-caudal axis (**Fig. S7A**). Model predictions were consistent with actual pathology readouts for regions that have injections reported, including the olfactory bulb (MOB), cortex (MOs), hippocampus (CA1/CA3/DG), substantia nigra (SN), and pedunculopontine nucleus (PPN)^43–46^. Additional injection sites include the medial septum (MS/NDB), thalamus (VAL), and dentate nucleus of the cerebellum (DN). Although the model predicts pathology patterns for these sites that are consistent with connectivity-mediated spread, experimental validation will be required to determine whether pathology develops as predicted. For example, thalamic nuclei express low α-synuclein and therefore may be resistant to developing pathology.

Turning back to predictions from the caudoputamen injection site, we hypothesized that pathology progression is influenced not only by anatomical connectivity but also by cell- and region-intrinsic factors such as cell type and gene expression. This differential regional vulnerability becomes apparent when model predictivity is visualized by anatomical division, with regions color-coded by anatomical group (**Fig. S7B**). Thalamic and mesencephalic regions exhibit lower pathology than predicted by the model. Assuming that local pathology results from a combination of anatomical connectivity and vulnerability factors, we estimated regional vulnerability as the difference between predicted and observed pathology (i.e., the residuals from the model fit to pathology data). Residuals were calculated for each region at each time point, and Pearson’s correlation coefficients were used to assess their similarity across hemispheres and time points (**Fig. S7C**). Overall, regional residuals were highly consistent across hemispheres, though less stable at early time points (0.1-0.5 MPI), likely due to the lower levels of pathology available to fit the model at those time points. Given the high conservation of residuals across hemispheres and from 1 to 9 MPI, we averaged those values to generate a composite regional vulnerability measure for subsequent analyses.

### Spatial neuronal gene expression atlas (PANGEA)

We hypothesized that the residuals of each region in the network diffusion model (i.e., the variance not explained by connectivity) are explained, at least in part, by regional gene expression. To test this hypothesis, we aimed to directly compare the unexplained variance to regional gene expression. To accomplish this, we needed a spatially resolved gene expression brain atlas. The first, and still one of the most widely used of these atlases is the ABA *in situ* hybridization data^47^. This massive resource is excellent for gene expression visualization but was not generated with the goal of regional gene quantification. While we and others have previously used this resource, only 4277 genes have expression patterns available after applying a standard quality control threshold^29,48^, and spatial registration is done using an automated pipeline and not validated for accurate anatomical annotation. A recent spatial transcriptomics atlas acquired with evenly-spaced spots covers a similar number of genes, but with quantitative infrastructure^49^. Other spatial transcriptomics technologies like MERFISH have enabled high-resolution transcript assignment but only cover hundreds of genes^50^. Each of these atlases provides valuable insight into molecular classification of cells and anatomical regions. However, to optimally determine the relationship between gene expression and neuronal vulnerability to pathology, we required regional expression, quantitative whole transcriptome coverage, and neuronal expression specificity.

We therefore developed the sPAtial Neuron Gene Expression Atlas (PANGEA) by performing GeoMx whole transcriptome atlas capture on segmented neurons in spatially registered brain sections (**Fig. 4A, 4B**). Coronal brain sections from 3 male and 4 female mice were stained with NeuN/HuC/HuD to label neuronal cell bodies and glial fibrillary acidic protein (GFAP), which served as a morphological marker for anatomical registration due to its strong signal in white matter tracts. Brain sections were registered to the ABA CCFv3 using the QUINT workflow, and registrations were imported onto the GeoMx instrument to enable anatomically guided region-of-interest designation (**Fig. 4C**). The area of illumination for barcode collection and sequencing was based on segmentation of NeuN/HuC/HuD immunofluorescence. Thus, transcript barcodes covering the whole transcriptome were captured from neuronal soma in anatomically defined regions. Altogether, the atlas represents 302 of the 722 gray matter regions in the ABA CCFv3. Regions that were not sampled include those corresponding to cortical layer 1, which has few neurons, and hindbrain regions that are difficult to anatomically distinguish accurately. After quality control, the atlas covers the regional expression of 19,781 genes.

To discern general trends in the data, we performed a principal component analysis on all regions (**Fig. 4D**). Regions clustered within principal components 1 and 2 based on major brain divisions, with cortical regions separating from subcortical regions. To further examine the utility of this atlas, we examined the expression of known cell type and region markers, and then we compared these data to the ABA *in situ* hybridization atlas, our previous *Snca* atlas^51^, and RiboTRAP data^52^ (**Fig. 4E, 4F**, **S8**). We found that the excitatory neuron marker *Slc17a7* is highly expressed in regions known to have excitatory projection neurons (cortex, basolateral amygdala), while the expression of inhibitory neuron marker *Gad2* was lower in these regions and more highly expressed in subcortical regions (**Fig. 4E**). Regional expression of these genes also correlated well with the “expression energy” of the same genes in the ABA *in situ* hybridization atlas. In contrast, the catecholaminergic neuron marker *Th* was largely restricted to the substantia nigra, ventral tegmental area, and locus coeruleus, as expected. The regional correlation of *Th* to the ABA was low, likely due to its sparse expression and difficulties with regional registration in the ABA expression atlas. Known cortical layer markers also showed highly stereotyped regional expression across the neuraxis—*Cux2* in layer 2/3, *Etv1* in layer 5, *Rprm* in layer 6 (**Fig. 4F**). *Snca*, the gene encoding α-synuclein, shows high expression in cortex, especially layer 5, amygdala, hippocampus and substantia nigra pars compacta, and low expression in many thalamic and mesencephalic nuclei (**Fig. S8A**). This is consistent with previous evaluation of *Snca* mRNA expression and α-synuclein protein expression that had been targeted to the nucleus (**Fig. S8B, S8C**)^51^.

To further establish the utility of this dataset, we examined the brain-wide correlation of gene expression to the 4277 genes available in the ABA dataset. We found that PANGEA and the ABA data were positively correlated for most genes (Pearson’s r mean=0.27641, minimum= - 0.50310, maximum= 0.92024) (**Fig. S8D**). Genes that were not positively correlated likely represent non-neuronal genes which were not sampled in our atlas, or reflect discrepancies in spatial registration, as with *Th*. To determine the depth to which region-selective gene expression could be characterized with this method, we evaluated the expression of the top 10 genes previously identified as dopaminergic neuron-selective using dopamine-driven RiboTRAP^52^ or as enriched in the substantia nigra relative to the ventral tegmental area. We found that all 10 genes (*Chrna6, Cpne7, Ddc, En1, Gch1, Pitx3, Slc6a3, Slc10a4, Slc18a2*, and *Th*) were strongly enriched in the substantia nigra pars compacta in PANGEA (**Fig. S8E**).

These examples highlight that PANGEA may be used not only to validate previously established markers, but also to understand anatomically-annotated full-genome neuronal transcription. PANGEA data can be browsed and downloaded at https://lume.tv/PANGEA/.

### Regional vulnerability is associated with specific gene expression pathways

The goal of generating PANGEA was to determine if there are gene expression patterns that are related to regional vulnerability (**Fig. 5A**). We therefore examined the correlation between the expression patterns of each gene and the regional vulnerability to α-synuclein pathology. As expected, *Snca* shows a positive correlation with vulnerability (**Fig. 5B**). However, *Snca* did not show the highest correlation with vulnerability or resilience of all genes, suggesting that there are gene expression patterns that more strongly influence whether a region develops α-synuclein pathology than the substrate gene itself. *Pld3* showed the highest correlation with vulnerability, while *Nefh* showed the most negative correlation (**Fig. 5B**).

**Figure 5.**
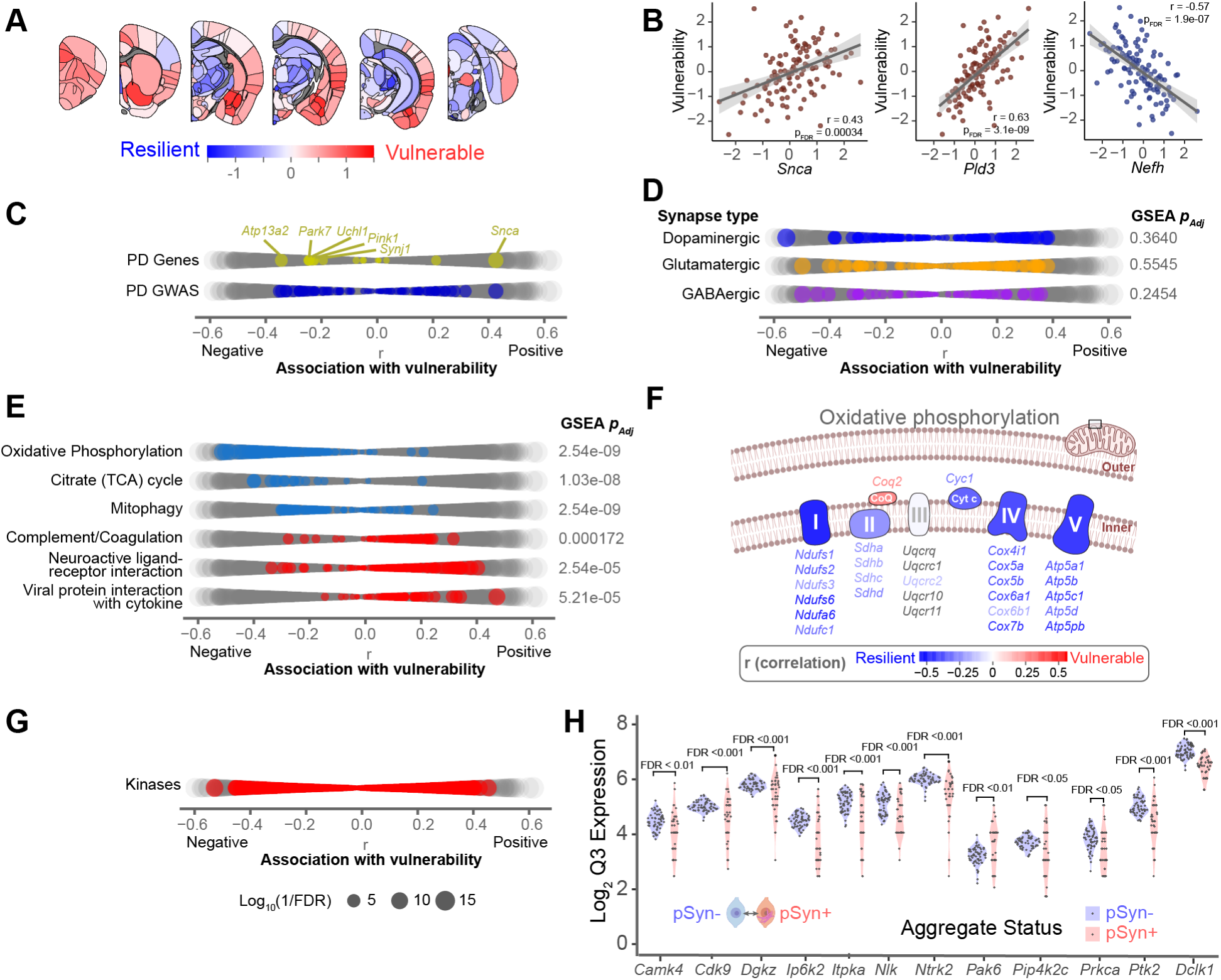
Regional vulnerability is associated with specific gene expression pathways. (A) Regional vulnerability heatmap plotting average region residuals (1,3,6,9 MPI) from the linear network diffusion model of pathology spread. (B) Scatterplot of *Snca* and top genes associated with vulnerability from PANGEA (Pearson’s *r*). Each dot represents the expression and vulnerability score (residual) from each region in PANGEA. Solid lines represent the line of best fit, and shading represents 95% confidence intervals. (C) Dot plot highlighting causal PD genes, and GWAS identified PD genes. Dots are scaled according to the significance of the correlation between gene expression and regional vulnerability (FDR corrected *p*-value). (D) Dot plot highlighting synaptic genes associated with specific neuron subtypes. (E) Dot plot highlighting genes of top gene sets identified as differentially enriched by GSEA. (F) Schematic of the oxidative phosphorylation electron transport chain. Genes relevant to each component of the electron transport chain are colored by their association to regional pathology vulnerability. (G) Dot plot highlighting kinase genes. (H) Violin plots of the gene expression level of 12 kinases positively associated with regional pathology vulnerability that also showed differential expression between neurons with and without α-synuclein inclusions in the cortex of the α-synuclein PFF model^40^.

Among causative and risk genes for PD, *Snca* showed the highest correlation with regional vulnerability (**Fig. 5B, 5C**). While *Snca* was the only PD causative gene with a significant positive correlation with vulnerability, several causative genes had a significant negative relationship with vulnerability (**Fig. 5B**). That is, they are more highly expressed in regions that are less likely to develop α-synuclein pathology. Mutations in all these genes (*Atp13a2, Park7, Uchl1, Pink1, Synj1*) are thought to lead to loss-of-function. The higher expression of these genes in regions that are less likely to develop aggregates suggests that their function may be protective in PD and that mutations, or lower expression, may lead to an increased propensity to develop pathology. None of the GWAS genes had a higher correlation with vulnerability than familial PD genes.

We next examined the relationship between vulnerability and known cell type markers, since excitatory and catecholaminergic neurons are thought to be more vulnerable to developing α-synuclein pathology (**Fig. 5D**). None of the gene sets relating to dopaminergic, excitatory, or inhibitory synapses were significantly enriched related to vulnerability. This could be due to the intrinsic mixture of cell types in many brain regions, or the redundancy of genes expressed in different synapse types.

We next aimed to identify gene pathways associated with vulnerability in an unbiased manner. We therefore performed gene set enrichment analysis to identify gene sets significantly associated, either positively or negatively, with vulnerability (**Fig. S9**). Overall, we found 70 gene sets that were significantly enriched for vulnerability or resilience to pathology (FDR<0.05). Mitochondrial oxidative phosphorylation, citrate cycle, and mitophagy were all highly over-represented in the resilient side (**Fig. 5E**, **S9**). Even at the individual gene level, oxidative phosphorylation genes are more highly expressed in regions that are more resilient to α-synuclein pathology (**Fig. 5F**). This increased expression could represent a conserved resilience pattern and regions with lower expression of these pathways intrinsically may be more susceptible to toxicity induced by mitochondrial perturbation, including by α-synuclein pathology^40^. Gene patterns positively related to vulnerability included the complement and coagulation cascade, neuroactive ligand-receptor interaction, and viral protein interaction with cytokine (**Fig. 5E**, **S9**). The complement and coagulation cascade is a key process in innate immunity, regulating the body’s response to foreign pathogens. The positive relationship between this cascade and vulnerability suggests higher regional expression of these genes relates to increased regional vulnerability to pathology.

While each of these associations provides evidence for pathways involved in the development of pathology in PD, many are difficult to manipulate due to their essential nature or lack of druggable targets. We therefore focused on a class of genes—kinases—that are amenable to manipulation to assess whether network-derived vulnerability is related to cellular vulnerability and can reveal novel therapeutic targets. Although kinases as a class showed no directional enrichment with respect to vulnerability, we postulated that kinases enriched in vulnerable regions could serve as effective targets for inhibition. We identified 106 kinases whose expression was significantly related to vulnerability (Pearson’s *r*, FDR<0.05) (**Fig. 6G**). To further narrow this list, we only included genes that we had previously identified as significantly different between α-synuclein inclusion-bearing neurons and their neighbors^40^. The final list included 12 kinases (*Cdk9, Nlk, Pak6, Dclk1, Prkca, Ptk2, Ntrk2, Ip6k2, Camk4, Itpka, Pip4k2c*, and *Dgkz*) that we hypothesized may have a direct relationship with regional neuronal vulnerability to developing α-synuclein pathology (**Fig. 5H**).

**Figure 6.**
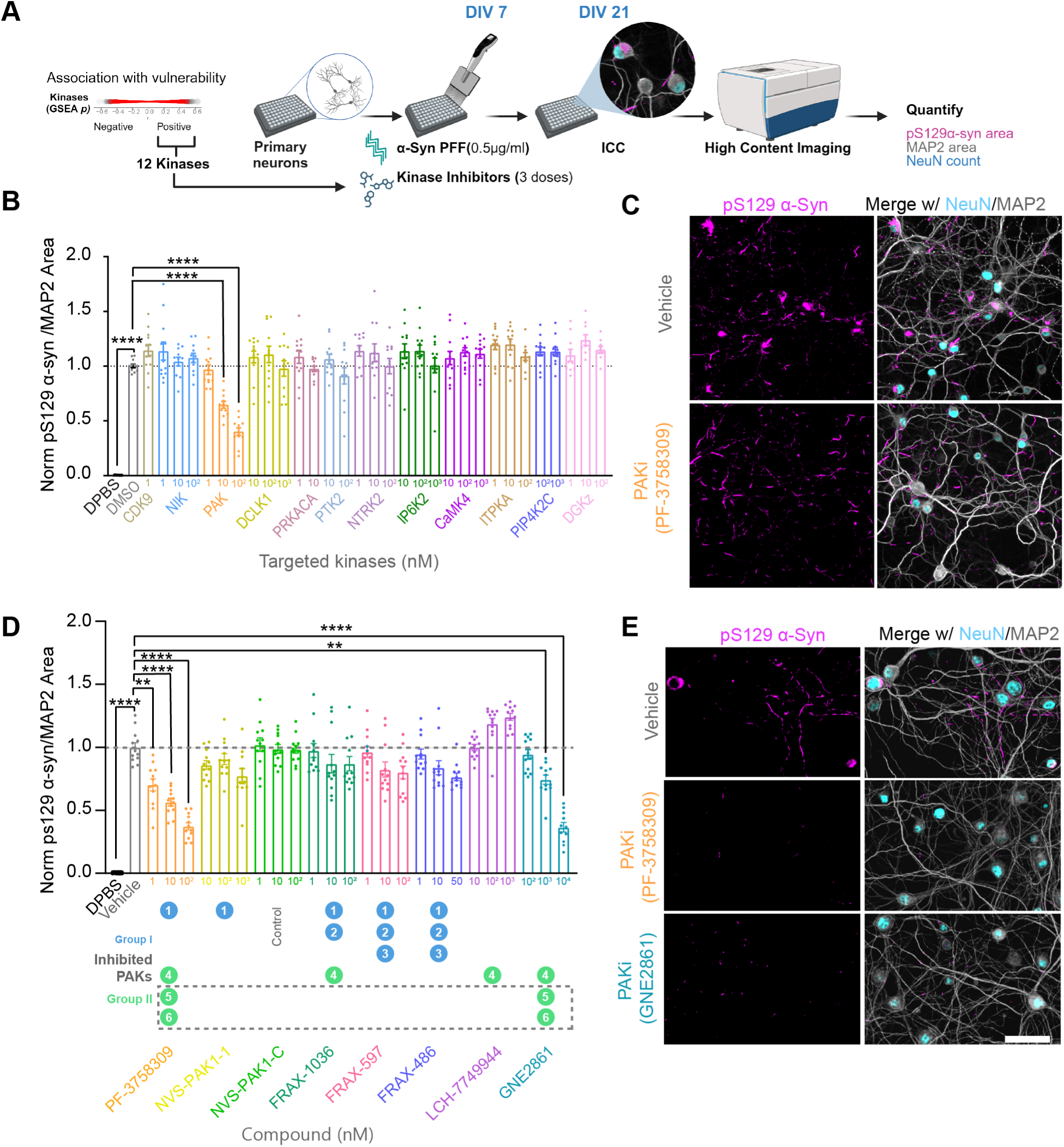
PAK inhibition is protective in a neuronal model of α-synucleinopathy. (A) Schematic for kinase inhibitor screen. (B) Kinases were inhibited at 3 different doses in primary hippocampal neurons treated with α-synuclein PFFs and pathology was quantified relative to MAP2 area. Data presented as mean ± SEM with individual values plotted. N=12 independent wells from 4 separate cultures. *p*-values represent fold-change compared to vehicle control with Welch ANOVA test and Dunnett’s T3 multiple comparison test: \*\*\*\**p* < 0.0001. (C) Representative image of pS129 α-synuclein in vehicle treated neurons compared to neurons treated with 100 nM of PAK inhibitor. (D) PAK inhibitors with different selectivity were used at 3 different doses in primary hippocampal neurons treated with α-synuclein PFFs and pathology was quantified relative to MAP2 area. Data presented as mean ± SEM with individual values plotted. N=12 independent wells from 4 separate cultures. *p*-values represent fold-change compared to vehicle control with Welch ANOVA test and Dunnett’s T3 multiple comparison test: **p < 0.001, ****p < 0.0001. (E) Representative images of pS129 α-synuclein in vehicle treated neurons compared to neurons treated with the highest dose of PF-3758309 or GNE2861. Scale bars = 50 µm.

### PAK inhibition is protective in a neuronal model of α-synucleinopathy

To evaluate the role of these 12 kinases at the cellular level, we performed a kinase inhibitor screen in a primary neuron model of α-synucleinopathy. Primary hippocampal neurons were cultured and transduced with α-synuclein PFF at 7 days *in vitro* (DIV) to induce pathology (**Fig. 6A**). On the same day, neurons were treated with kinase inhibitors or vehicle control. Neurons were treated with kinase inhibitors at three doses spanning values above and below their reported IC50. Neurons were fixed 14 days after treatment and stained for NeuN, MAP2, and pS129 α-synuclein to evaluate the effect of the test compounds on neuronal health and α-synuclein pathology. Some compounds induced neuron death (**Fig. S10A, S10B**), which could confound measurements of pathology. Therefore, treatments which caused more than a 25% reduction in NeuN count or MAP2 area were omitted from further analysis. Pathology was quantified as the pS129 α-synuclein positive area normalized to MAP2 area. The PAK inhibitor PF-3758309 induced a dose-dependent reduction in α-synuclein pathology, with the maximum tested dose of 100 nM reducing pathology by around 50%. **(Fig. 6B, 6C)**. No other compound tested in this initial screen of 12 kinases significantly reduced α-synuclein pathology (**Fig. 6B, S10**).

The PAK family is composed of six members, which are categorized into two groups: Group I (PAK1, PAK2, and PAK3) and Group II (PAK4, PAK5, and PAK6). Since PF-3758309 inhibits PAK1 from group I and PAKs 4,5 and 6 from group II^53^, we aimed to identify the specific PAKs contributing to α-synuclein pathology. To do this, we tested a suite of PAK inhibitors with differential specificity across the PAK family (**Fig. 6D, 6E**). We did not observe a significant reduction in α-synuclein pathology with NVS-PAK1-1, a highly specific PAK1 inhibitor, as compared to its control, NVS-PAK1-C. Similarly, other PAK inhibitors like FRAX-1036 (inhibiting PAK1,2,4), FRAX-597 (inhibiting PAK1,2,3), and FRAX-486 (inhibiting PAK1,2,3) failed to reduce α-synuclein pathology. We saw a trend toward increased α-synuclein pathology with LCH-7749944, a PAK4 specific inhibitor, which is consistent with a previous study^54^. One additional compound, GNE2861, which is specific for Group II PAKs, induced a dose-dependent reduction in pathology, with approximately 50% reduction observed at the highest tested dose of 10 μM (**Fig. 6D**, **6E**). Given that PF-3758309 (inhibits PAK1, 4, 5, and 6) and GNE-2861 (inhibits PAK4-6) reduced α-synuclein pathology, whereas PAK1 and PAK4 inhibitors did not, inhibition of PAK5 and/or PAK6 is likely responsible for the observed reduction in pathology.

The doses at which PAK inhibitors were tested were well tolerated and did not induce significant cell death (**Fig S10C, S10D**). We moved forward with GNE2861 due to its increased specificity for group II PAKS, in contrast to PF-3758309, which also inhibits PAK1.

### Group II PAK inhibition or genetic depletion is neuroprotective even with delayed treatment

Although PAKs have not been reported to phosphorylate α-synuclein, we wanted to verify that the reduction in pS129 α-synuclein signal observed following group II PAK inhibition was indeed a reduction in α-synuclein pathology and not simply a reduction in phosphorylated α-synuclein. To test this possibility, neurons were treated as before with GNE2861 concurrently with human α-synuclein PFFs. Fourteen days later, soluble proteins were extracted with 1% TX-100 during fixation, and neurons were stained for total α-synuclein and pS129 α-synuclein antibody (**Fig. 7A**). Both insoluble total and pS129 α-synuclein were reduced by GNE2861 (**Fig. 7B-7D**), further demonstrating that PAK inhibition decreases the accumulation of α-synuclein pathology.

**Figure 7.**
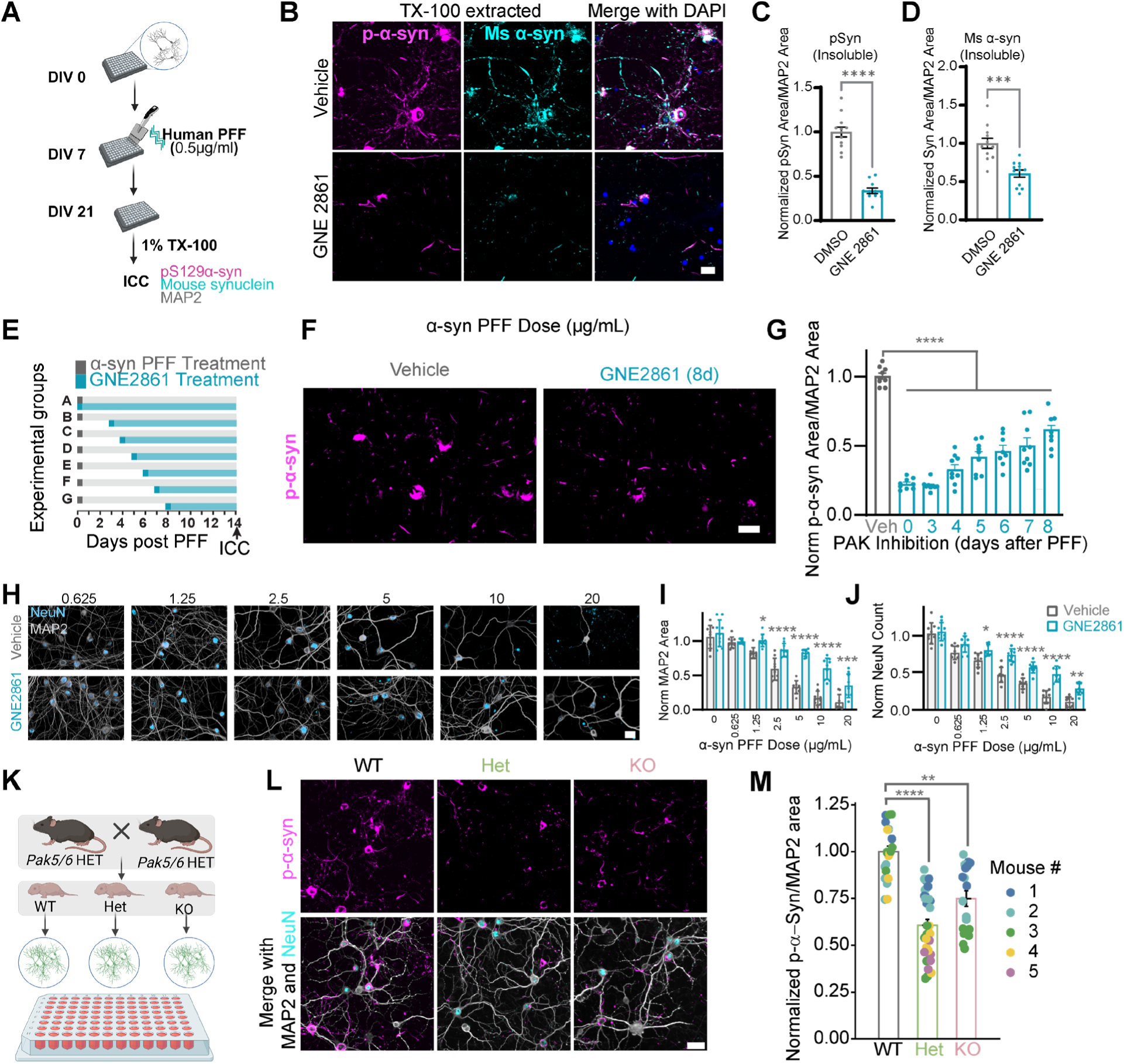
Group II PAK inhibition or genetic depletion is neuroprotective even with delayed treatment. (A) Experimental schematic for soluble protein extraction during immunostaining. (B) Representative images of insoluble pS129 α-synuclein and total mouse α-synuclein after treatment of vehicle or 10 μM GNE2861. Scale bar = 10 µm. Both (C) insoluble pS129 α-synuclein (D) and insoluble total mouse α-synuclein were reduced on GNE2861 treatment. N=9 independent wells from 3 separate cultures, statistical significance was calculated using an unpaired two-tailed t-test with Welch’s correction, ***p < 0.001, ****p < 0.0001. (E) Schematic of the delayed treatment of neurons with PAK inhibitor following α-synuclein PFF treatment. (F) Representative images of pS129 α-synuclein pathology after 8-day delayed treatment of 10 μM GNE2861. (G) Quantification of α-synuclein pathology in neurons following GNE2861 treatment at different intervals after α-synuclein PFF treatment. N=9 independent wells from 3 separate cultures. *p*-values represent fold-change compared to vehicle control with Welch ANOVA test and Dunnett’s T3 multiple comparison test: ****p < 0.0001. (H) Representative images of primary neurons treated with increasing doses of α-synuclein PFFs. Both somatodendritic marker MAP2 (I) and neuronal nuclei marker NeuN (J) show dose-dependent reductions and are rescued by treatment with GNE2861. N=9 independent wells from 3 separate cultures. Data presented as mean ± SEM with individual values plotted. *p*-values represent fold-change compared to vehicle control with 2-way ANOVA test with Sidak’s multiple comparison test, *p <0.05, ***p <0.01, ***p = 0.001,****p < 0.0001. (K) Schematic of the genetic validation strategy, heterozygous *Pak5/Pak6* knockout mice were bred, and hippocampal neurons from individual pups were cultured separately to generate wild-type (WT), heterozygous (Het), and knockout (KO) groups. Tail samples from each pup were used to confirm genotypes. (L) Representative images of α-synuclein pathology in primary neurons. (M) Quantification of α-synuclein pathology in primary neurons from each genotype following treatment with α-synuclein preformed fibrils (PFFs). Data presented as mean ± SEM with individual mice indicated by dot color. Statistical analysis was performed using a mixed-effects model, followed by Dunnett’s multiple comparisons test comparing each group to WT. N = 4 (WT), 5 (Het), and 3 (KO) independent cultures. Adjusted p-values: ****p < 0.0001 (WT vs. Het), **p = 0.0016 (WT vs. KO). Scale bars = 20 µm.

Several potential mechanisms could underlie the reduction of pathology and improvement in neuronal health, including decreased α-synuclein PFF internalization, inhibition of PFF escape from lysosomes, decreased α-synuclein monomer recruitment, and increased degradation of fully formed aggregates. Treating neurons with inhibitors and α-synuclein PFFs simultaneously does not enable us to distinguish between these possible mechanisms. To gain insight into how group II PAK inhibition reduces pathology and rescues neuron health, neurons were treated with the group II PAK inhibitor 0, 3, 4, 5, 6, 7, or 8 days after α-synuclein PFF treatment (**Fig. 7E**). Remarkably, GNE2861 treatment reduced α-synuclein pathology even when administered up to 8 days after PFF treatment (**Fig. 7F, 7G**). α-Synuclein PFFs are fully internalized within 24 hours of treatment^55^, so the fact that treatment with inhibitor reduces pathology even 8 days after α-synuclein PFF administration suggests that group II PAK inhibition does not act on the initial steps of internalization or lysosomal escape, but rather on cytosolic aggregate formation or clearance.

In primary neurons, the α-synuclein PFF dose previously used (0.5 μg/mL) does not induce neuronal death over 21 DIV. We therefore treated neurons with increasing doses of α-synuclein PFFs (0.625 to 20 ug/mL) and aged neurons to 28 DIV. This treatment paradigm induced a significant dose-dependent reduction of MAP2 area (**Fig. 7H, 7I**) and neuron count (**Fig. 7H, 7J**). Treating neurons with 10 μM of GNE2861 rescued the reduction of NeuN count and MAP2 area at each tested dose (**Fig. 7H-7J**).

To assess off-target effects at the tested concentrations, we performed a follow-up experiment using a lower dose of PFFs (0.125 μg/mL) to reduce pathology burden and evaluated compound efficacy across a reduced concentration range of GNE2861 (**Fig. S12E-S12H**). GNE2861 decreased α-synuclein pathology at 500 nM, with significant effects observed at 1 μM and maximal suppression at 2 μM, beyond which no further reduction was observed (**Fig. S12E-S12H**). The retention of efficacy of GNE2861 at lower concentrations, together with efficacy of a second inhibitor (PF-3758309), reduces the likelihood that these effects are driven by off-target mechanisms. Consistent with target engagement, GNE2861 reduced phosphorylated group II PAK (pPAK) levels in primary neurons under both PBS and PFF conditions compared to vehicle (**Fig. S12I-S12J**).

To further validate group II PAKs as modulators of α-synucleinopathy, we next employed a genetic approach. Primary neurons were cultured from pups derived from a heterozygous *Pak5/6* knockout (KO) cross. Because *Pak5* and *Pak6* are located on the same chromosome, they segregate together, yielding wildtype, heterozygous, or homozygous KO animals for both genes (**Fig. 7K**). Neurons from individual mice were treated with α-synuclein PFFs at DIV 7, and pathology was evaluated 14 days later. Both heterozygous and homozygous KO neurons exhibited a significant reduction in α-synuclein pathology relative to WT (**Fig. 7K-7M**). Notably, the magnitude of reduction was similar between heterozygous and homozygous neurons, indicating that partial depletion of PAK5/6 is sufficient to suppress aggregation without affecting neuronal health (**Fig. S12K-12L**). Together, these pharmacological and genetic results further support a role for PAK5/6 in regulating α-synuclein pathology.

### Group II PAK inhibition reduces α-synuclein aggregation *in vivo*

A previous study found that PAKs phosphorylated at an activating residue were elevated in PD brains^56^ and our previous data suggest increased PAK6 expression in pSyn inclusion-bearing neurons both in PFF injected mice and human PD brain^40^ (**Fig. S12A-S12D**). To further define the relationship between PAK activation and α-synuclein pathology, we stained primary neurons, mouse brain tissue, and human brain tissue for α-synuclein inclusions and phosphorylated PAK (pPAK), detected with an antibody that recognizes an autophosphorylation site on group II PAKs^57^ (**Fig. 8A-G**). In primary neurons, 81% of inclusion-bearing neurons contained pPAK puncta compared to 4% of neurons without inclusions (**Fig. 8A-8B, S11**). A similar pattern was observed *in vivo,* with pPAK present in 87% of inclusion-bearing neurons of 3 MPI mice, compared to 0.4% of inclusion-negative neurons (**Fig. 8C-8D**). Dopaminergic neurons in the substantia nigra showed a lower proportion of pPAK positivity (69% positive, **Fig. 8E-8F**), suggesting cell type-specific differences in PAK activation. In the human PD substantia nigra pPAK puncta were also observed in a subset of Lewy body-bearing neurons (**Fig. 8G**); however, limited pathology prevented robust quantification. No punctate pPAK signal was detected in α-synuclein monomer-injected mice (**Fig. S11C**), and similar localization was observed using an independent pPAK antibody (**Fig. S12M**). pPAK signal was also significantly reduced in Pak5/6-deficient primary neurons, although residual staining remained (**Fig. S12N-S12O**), suggesting that the antibody detects pPAK5/6-dependent signal but also exhibits some non-specific binding.

**Figure 8.**
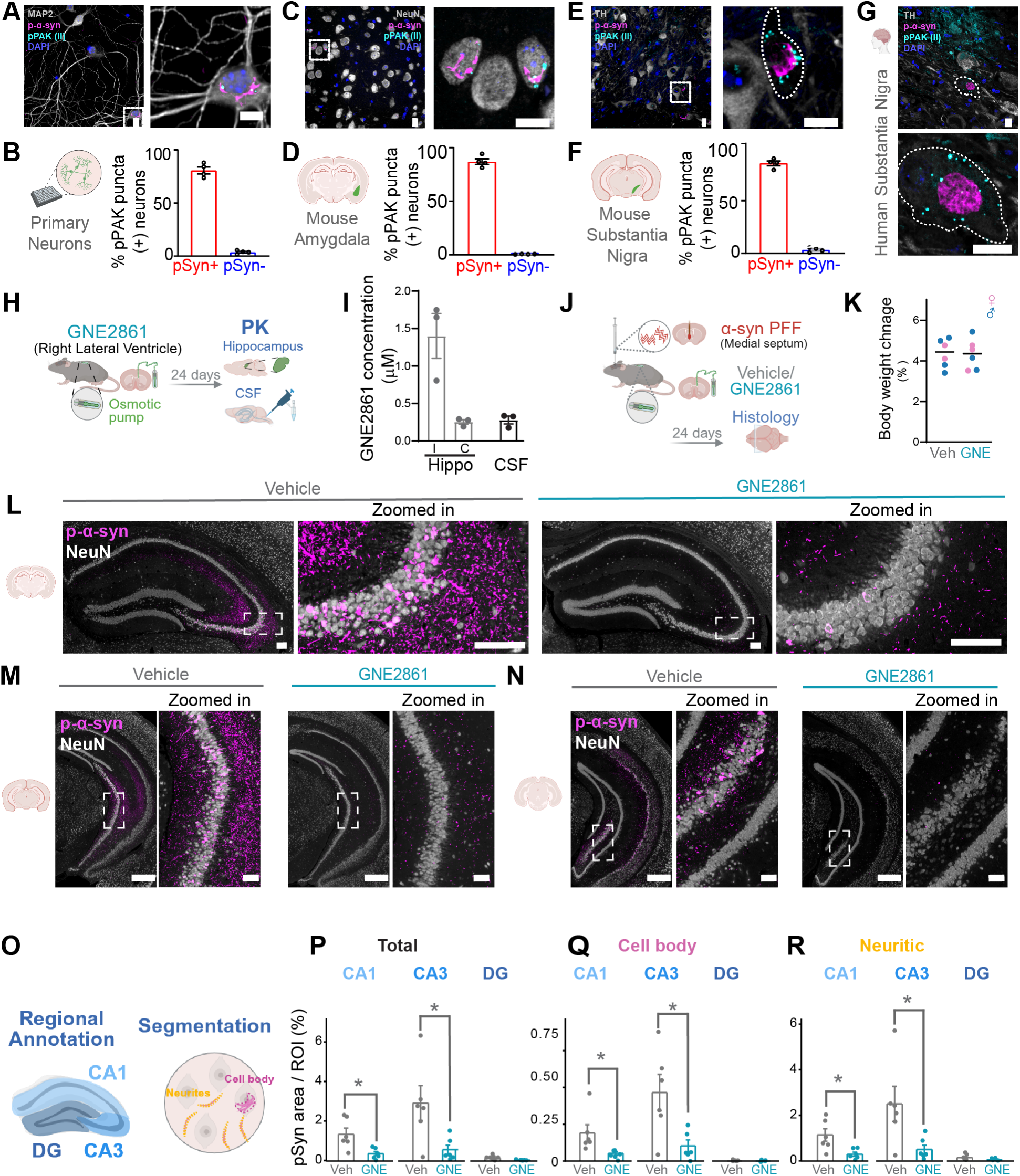
Group II PAK inhibition reduces α-synuclein aggregation *in vivo*. Representative images for co-immunofluorescence of α-synuclein inclusions and pPAK (group II) that shows punctate distribution of PAKs near inclusions in (A) primary neurons, (C) mouse amygdala, (E) mouse substantia nigra, and (G) human PD substantia nigra. TH = tyrosine hydroxylase. Scale bars = 10 μm. Quantification of the percentage of neurons that were pPAK-positive in neurons with or without pS129 α-synuclein inclusions in (B) primary neurons, N=12 independent wells from 4 separate cultures, (D) mouse amygdala, N=4, and (F) mouse substantia nigra, N=4. (H) Schematic of the experimental design for the pharmacokinetic (PK) study using intracerebroventricular (ICV) infusion of GNE2861 in the right lateral ventricle via ALZET osmotic minipump. (I) PK analysis of GNE2861 in the hippocampus (ipsilateral and contralateral to infusion) and cerebrospinal fluid (CSF) collected 24 days post-surgery, demonstrating levels of the compound. (J) Schematic of the experimental design for the pharmacodynamic study (PD) study using intracerebroventricular (ICV) infusion of GNE2861 in the right lateral ventricle via ALZET osmotic minipump and α-synuclein PFF in the medial septum. (K) Body weight assessment from the PD cohort. Percent change in body weight was calculated relative to post-surgery baseline and measured at the study endpoint (24 days post-surgery) in vehicle- and GNE2861-treated mice. Sections from rostral, mid, and caudal hippocampus were analyzed to capture the full rostrocaudal extent. (L–N) Representative hippocampal sections are shown for vehicle- and GNE2861-treated mice injected with α-synuclein PFFs injected. Scale bars = 100 μm. (O) Schematic of regional annotation and segmentation. Hippocampal subregions (DG, CA1, CA3) were defined for quantification, and within each region, inclusion burden was measured as total, cell body, and neuritic. (P) Total, (Q) cell body, and (R) neuritic pS129 α-synuclein burden was quantified for each of the 3 regions. Data represent mean percentage area occupied/ROI area per mouse (mean of left and right hemispheres). N = 6 mice per group. Statistical significance was calculated using an unpaired two-tailed t-test with Welch’s correction, *p < 0.05.

To determine whether pharmacological inhibition of PAKs is effective *in vivo*, we first assessed the brain penetrance of GNE2861. *In silico* analysis using SwissADME predicted poor blood-brain barrier permeability^58^, which was confirmed experimentally following intravenous administration. To overcome this limitation, GNE2861 was administered through chronic intracerebroventricular (ICV) infusion into the right lateral ventricle using an ALZET osmotic pump (**Fig. 8H**). Because ICV delivery results in higher exposure near the ventricular system, the hippocampus was selected a priori for assessment based on its proximity to the lateral ventricle (**Fig. S12P**) and previous research demonstrating optimal exposure there following ICV administration^59^. For pharmacokinetic (PK) analysis, pumps loaded with 6 mM GNE2861 were implanted in mice, and cerebrospinal fluid (CSF) and hippocampi were collected after 24 days. GNE2861 concentrations reached approximately 0.28 µM in the CSF and contralateral hippocampus and approximately 1.4 µM in the ipsilateral hippocampus (**Fig. 8I**), within the range of effective *in vitro* doses.

To achieve robust hippocampal pathology, α-synuclein PFFs were injected into the medial septum, which drives bilateral α-synuclein pathology in the hippocampus^60^. Osmotic pumps were implanted immediately after α-synuclein PFF injection (**Fig. 8J**). Treatment was well tolerated for the 24-day study, as indicated by normal weight gain in mice (**Fig. 8K**) and the absence of any overt phenotypes. Residual compound recovered from pumps at the end of the study retained protective activity in primary neurons, confirming compound stability of GNE2861 during *in vivo* delivery (**Fig. S12Q-S12U**).

To assess the impact of PAK inhibition on α-synuclein pathology, brain sections corresponding to hippocampal levels (ABA CCFv3 sections 72, 82, and 88) were collected and stained for pS129 α-synuclein and NeuN. GNE2861 treatment resulted in a marked reduction in α-synuclein aggregation across hippocampal regions (**Fig. 8L-8N**). Quantification across hippocampal regions (CA1, CA3, and dentate gyrus) (**Fig 8O, S12V-S12X**) revealed a consistent 75% or greater reduction in total, somatic, and neuritic pathology (**Fig. 8P-8R**). To assess target engagement, hippocampal sections were stained for phosphorylated group II PAK (pPAK). GNE2861 treatment resulted in an overall reduction in pPAK across hippocampal regions **(Fig. S12Y-S12Z)**.

## DISCUSSION

In this study, we sought to bridge the gap between network-level and cellular vulnerability to α-synuclein pathology. We combined high spatiotemporal resolution brain-wide pathology mapping, network diffusion modeling, and regionally resolved neuronal gene expression (PANGEA) to identify molecular determinants of vulnerability. The comparison of gene expression to regional vulnerability enabled the identification and validation of group II PAKs as potential therapeutic targets for PD. Given the lack of disease-modifying therapies for PD, these findings support a network-to-cell framework as a strategy for identifying therapeutic targets.

Our pathology mapping provides a detailed view of α-synuclein pathology across the mouse brain. Compared to prior studies^30,31,38^, this work extends spatial, temporal, and subcellular pathology resolution by quantifying neuritic and cell body inclusions across 8 timepoints and 1046 brain regions. In comparing three- and two-dimensional datasets, we found that they were largely correlated, with minimal discrepancies attributable to difficulties with accurate spatial registration and non-specific ventricular staining in three-dimensional datasets, and the potential for missed neuron populations in two-dimensional sampling. Pathology develops rapidly following α-synuclein PFF injection, with widespread pathology deposition evident by 9 days, initially localized ipsilaterally and spreading bilaterally over time. Regional differences in temporal dynamics were observed, with caudal cortical regions reaching peak pathology earlier than frontal cortical and subcortical regions. A key observation is that neuritic pathology precedes cell body pathology and remains the dominant form even at later timepoints when cell body pathology has matured. This shift from neuritic pathology to cell body pathology is consistent with the shift in aggregate size distribution described by Dadgar-Kiani and colleagues^38^. These findings suggest that relative proportions of neuritic and cell body pathology may provide a useful proxy for the stage of pathology progression, with implications for interpretation of human neuropathology.

To place these regional and cellular patterns in a broader systems-level context, we applied a linear network diffusion model. Network diffusion modeling has previously been used to predict the spread of α-synuclein and tau^29–37^ and to identify genetic and cellular factors associated with regional vulnerability^31,33–36^. These prior studies have generally attributed large-scale pathology patterns to network-driven progression through the structural connectome and have distinguished network-aligned and network-dissociated gene expression patterns^35^ as well as cell type composition associated with vulnerability to pathology^34,36^. Here, we extend this in high-resolution quantitative pathology datasets. Consistent with prior work^29–37^, we find that pathology progression is largely explained by retrograde connectivity, with modest improvement from incorporating bidirectional transport. The model performed better at predicting neuritic pathology than cell body pathology, likely reflecting earlier and broader distribution of neuritic aggregates, which increases dynamic range and statistical power for model fitting.

As more imaging ligands become available for assessing pathology or surrogate measures in human brain, computational modeling is becoming increasingly important for understanding disease progression. Recent studies modeling tau and amyloid β enabled by PET imaging ligands^17^ demonstrated that intracellular tau pathology appears to progress through anatomical networks, while amyloid β pathology appears to be more strongly associated with spatial proximity. In humans, computational modeling has been used to predict the site of pathology initiation, identify subtypes of disease, develop individualized pathological prognosis^20^, and identify regional gene expression patterns related to pathology progression patterns^15,18^. Given this human evidence that anatomical connectivity and gene expression networks are two major constraints on pathology progression, we calculated a regional vulnerability measure to compare to regional gene expression.

To enable this analysis, we developed PANGEA, a brain-wide atlas of neuronal gene expression covering 302 gray matter regions. While existing atlases^47,49,50,61–64^ are valuable resources, none offered the combination of anatomical resolution and neuronal specificity required for this study. PANGEA recapitulates known gene expression patterns and provides a platform for identifying novel spatial relationships. To make this dataset as accessible as possible, we developed an interactive web browser to display, analyze, and download the data: https://lume.tv/PANGEA/. The quality of the PANGEA dataset is reflected in the plotting of transcripts selective for dopamine neurons^52,65–67^. The power of PANGEA is its broad transcriptome coverage, and future integration with single-cell and cell type-resolved datasets will be valuable to refine vulnerability analysis.

Comparison of regional vulnerability with gene expression revealed associations with specific molecular pathways. As expected, *Snca*, the gene encoding α-synuclein, was positively correlated with vulnerability; however, several loss-of-function PD genes were significantly anti-correlated with vulnerability (*Atp13a2, Park7, Pink1, Synj1*), suggesting that lower gene expression in the absence of mutations may increase vulnerability to developing pathology. The pathways most negatively correlated with vulnerability were related to mitochondrial metabolism (oxidative phosphorylation, TCA cycle, mitophagy), suggesting that baseline metabolic capacity may be protective in α-synucleinopathies. This finding is consistent with the known role of mitochondrial dysfunction in PD^68^ and with genetic evidence implicating mitochondrial genes *PARK7* and *PINK1*^69^. In contrast, complement and cytokine pathways were positively correlated with vulnerability, consistent with prior observations of complement/coagulation cascade activation in A53T α-synuclein transgenic mice^70^, α-synuclein PFF-injected rats^71^, α-synuclein PFF-injected rats^71^, and PFF-injected mice^40,70^, as well as the brains of people with PD^72–75^. A recent study demonstrated that α-synuclein aggregates lead to complement C3 activation in astrocytes and enhanced C3 receptor activation and apoptosis in nearby neurons^76^. These findings suggest that baseline inflammatory tone may influence susceptibility to α-synuclein pathology.

Li et al.^36^ recently used a similar strategy to identify genes whose expression pattern is related to regional pathology development, albeit using 3855 genes from the Allen gene expression atlas^47^ and a model by which genes were incorporated into incoming or outgoing network edges to determine which approach improved model predictivity. Interestingly, despite the different datasets and modeling methods, there was some conservation of the overall gene sets identified as related to pathology progression (synaptic pathways) and even individual genes. While the top genes identified by Li et al. for individual pathways (*Dok5* and *Grid2ip*) were not significantly related to vulnerability in our analysis, *Pde1a*, the gene most strongly associated with their combined score, showed a positive association with vulnerability in our analysis (r=0.363, *p*_FDR_= 0.004). In addition, *Rora*, the gene whose overexpression was protective in a neuronal model of α-synucleinopathy in the Li et al. study, was associated with resilience in our dataset (r=-0.323, *p*_FDR_= 0.0139). Together, these results suggest convergence between network-level molecular predictors of pathology progression and cellular mechanisms of neuronal vulnerability and resilience.

While mitochondrial metabolism and complement/cytokine pathways are biologically informative, they are challenging to target therapeutically. Therefore, we focused on kinases that were positively associated with vulnerability and differentially expressed in neurons bearing α-synuclein inclusions^40^ as their inhibition might be expected to be protective. Of the selected 12 kinases, doublecortin-like kinase 1 (DCLK1) has been shown to regulate α-synuclein levels^77^, inositol hexaphosphate kinase 2 (IP6K2) is reported to be neuroprotective^78^, and PAKs have been associated with PD in several contexts^40,54,56,79^. Of 12 kinase inhibitors tested, only PAK inhibition produced a clear, protective effect on neurons. This effect was confirmed using a second, group II-selective inhibitor and by genetic depletion of PAK5/6, and was further validated *in vivo*, supporting group II PAKs as potential therapeutic targets in PD.

PAKs are serine/threonine kinases involved in cytoskeletal dynamics, vesicle trafficking, and signal transduction^80–83^. Group II PAKs (PAK4-6) have distinct roles from group I PAKs, including regulation of presynaptic vesicle dynamics^84–86^, and show enriched expression in the brain^87–89^. PAK6, in particular, is co-expressed with androgen and glucocorticoid receptors in dopaminergic neurons^90^, suggesting a role in this vulnerable cell population in PD. Despite broad brain expression of PAK5 and PAK6^87–89^, KO for either or both of these genes in mice leads to viable, fertile progeny with no morphological defects^87^. In contrast, *Pak4* KO leads to embryonic lethality due to cardiac abnormalities. PAK4 has been shown to limit α-synuclein aggregation in overexpression-based systems^54,79^. Similarly, in our screen, we found slight increases in pathology with a selective PAK4 inhibitor. Together with our findings from PAK5/6 genetic depletion, this suggests that protection against α-synuclein pathology is conferred by inhibition of PAK5 and PAK6, but not PAK4.

PAK6 has been shown to interact with PD protein leucine-rich repeat kinase 2 (LRRK2)^56^. Group II PAKs phosphorylated at an activating residue are elevated in both idiopathic and *LRRK2*-PD, while total PAK6 is only elevated in *LRRK2*-PD^56^. While we find an elevation of PAK6 transcript in inclusion-bearing neurons, together these data suggest that post-translational modification of PAK is a primary means of modulation of PAK kinase activity in idiopathic PD. PAK6 was found to rescue LRRK2^G2019S^-dependent changes in ciliogenesis^91^, but the localization of PAKs in our study did not appear at cilia, suggestive of a distinct mechanism. Instead, staining in primary neurons, mouse brain tissue, and PD brain tissue demonstrated that phosphorylation of group II PAKs was elevated almost exclusively in neurons surrounding inclusions, demonstrating the cell-autonomous nature of PAK activation. The punctate localization indicates that PAKs have accumulated in some type of signaling hub, possibly an organelle near α-synuclein inclusions. Interestingly, similar puncta were observed in AD brains stained for group I but not group II PAKs^92^, suggesting a mechanistic division between the two diseases. Further investigation is needed to determine the nature of these puncta, their potential localization within specific soma structures, and their role in α-synucleinopathy.

This study has several limitations. One limitation is the resolution of the connectivity data, and by extension the computational modeling and gene expression relationships. Current whole-brain connectivity maps are only available at mesoscale resolution, so cortical layers are not resolved. To facilitate future analyses at finer anatomical scales, we have collected pathology and gene expression data at full regional resolution, anticipating the development of anatomical connectivity maps with cortical laminar resolution. The limitations of coronal sampling compared to full three-dimensional reconstruction are discussed above. For the current study, this form of sampling is optimal, but the comparison of two- and three-dimensional data in the future may enhance knowledge of pathology progression. The computational model used in this study is also intentionally simple, relying solely on anatomical connectivity to predict patterns of pathology spread. This simplicity is a strength: it highlights the predictive power of the structural connectome alone and makes the model highly interpretable and computationally efficient. Its linear formulation provides a robust baseline for inference, with reduced flexibility and thus reduced susceptibility to overfitting compared to more complex models, and relative transparency in attributing spatial patterns to network structure versus local biological factors. However, the model’s simplicity also imposes limitations. Additional biological contributors to variation in regional pathology progression, such as neuron death, may be incorrectly interpreted as regional vulnerability. Future models incorporating non-linear spreading dynamics^13,16^ and other factors, such as production, clearance, aggregation rate and neuron death rate^13,36,37,93^ may improve predictivity and offer deeper insight into the mechanisms driving disease progression. Finally, our regional vulnerability maps are based on a single injection site. This injection site was chosen because mice develop pathology broadly in disease-relevant circuits such as the nigro-striatal and cortico-cortical circuits. However, generalizing this vulnerability map to alternative injection sites in the future would be beneficial. In addition, the mechanism by which PAK5/6 inhibition leads to a reduction in insoluble phosphorylated α-synuclein remains to be determined.

Overall, our study provides a proof-of-principle for the ability to translate network-level findings related to pathology progression to cellular targets for therapeutic intervention in PD. We also provide several resources for the neuroscience and neurodegeneration community to study pathology and gene expression patterns. Finally, we identify a neuroprotective class of kinase inhibitors for further therapeutic development in PD.

## MATERIALS AND METHODS

### Animals

All housing, breeding, and procedures were performed according to the NIH Guide for the Care and Use of Experimental Animals and approved by the Van Andel Institute Institutional Animal Care and Use Committee (IACUC). *Mus Musculus* CD1 (strain 022; RRID: IMSR_CRL:022) mice were obtained from Charles River and used for primary neuron culture. Wild type C57BL/6J mice (000664; RRID: IMSRJAX:000664) and Pak5/6 knock-out (KO) mice B6;129-*Pak6^tm1Amin^ Pak5^tm1Amin^*/J mice; RRID:IMSR_JAX:015825) were purchased from the Jackson Laboratory. Animals were kept on 12 h light/dark cycles, with food pellets and water *ad libitum*.

### α-Synuclein PFF

Purification of recombinant mouse α-synuclein and generation of α-synuclein PFFs was conducted as described elsewhere^94–96^. The pRK172 plasmid containing the gene of interest was transformed into BL21 (DE3) RIL-competent *E. coli* (Agilent Technologies Cat#230245). A single colony from this transformation was expanded in Terrific Broth (12 g/L of Bacto-tryptone, 24 g/L of yeast extract 4% (vol/vol) glycerol, 17 mM KH_2_PO_4,_ and 72 mM K_2_HPO_4_) with ampicillin. Bacterial pellets from the growth were sonicated and the sample was boiled to precipitate undesired proteins. The supernatant was dialyzed with 10 mM Tris, pH 7.6, 50 mM NaCl, 1 mM EDTA, and 1 mM phenylmethylsulfonyl fluoride (PMSF) overnight. Protein was filtered with a 0.22 µm filter and concentrated using Amicon Ultra-15 centrifugal filter units (Millipore Sigma Cat#UFC901008). Protein was then loaded onto a Cytiva HiLoad 26/600 Superdex 200 pg (Cytiva Cat#28989336) and 5 mL fractions were collected. Fractions were run on SDS-PAGE and stained with InstaBlue protein stain (ApexBio B8226) to select fractions that were highly enriched in α-synuclein. These fractions were combined and dialyzed in 10 mM Tris, pH 7.6, 50 mM NaCl, 1 mM EDTA, 1 mM PMSF overnight. Dialyzed fractions were applied to the MonoQ column (Cytiva, HiTrap Q HP 17115401) and run using a linear gradient from 25 mM NaCl to 1 M NaCl. Collected fractions were run on SDS-PAGE and stained with InstaBlue protein stain. Fractions that were highly enriched in α-synuclein were collected and dialyzed into DPBS. Protein was filtered through a 0.22 µm filter and concentrated to 7 mg/mL (α-synuclein) with Amicon Ultra-15 centrifugal filter units. Monomer was aliquoted and frozen at -80°C. For preparation of α-synuclein PFFs, α-synuclein monomer was shaken at 1,000 rpm at 37 °C for 7 days. Conversion to PFFs was validated by sedimentation at 100,000 x *g* for 60 minutes and by thioflavin S staining.

### Stereotaxic Injection

All surgery experiments were performed in accordance with protocols approved by the IACUC of Van Andel Institute. α-Synuclein PFFs were vortexed and diluted with DPBS to 2 mg/mL and sonicated in a cooled bath sonicator at 9°C (Diagenode Bioruptor® Pico; 10 cycles; setting medium; 30 seconds on, 30 seconds off). 3-4 month old mice were deeply anesthetized with isoflurane and injected unilaterally into the right forebrain targeting the dorsal striatum (coordinates: +0.2 mm relative to Bregma, +2.0 mm from midline, -2.5 mm beneath the dura) Injections were performed using a 10 µL syringe (Hamilton 7635-01, NV) with a 34-gauge needle (Hamilton 207434, NV) injecting 5 µg α-synuclein PFFs (2.5 µL) at a rate of 0.4 µL/minute. At the designated time points, mice were perfused transcardially with PBS and 4% paraformaldehyde (PFA), brains were removed and underwent overnight fixation in 4% PFA. After perfusion and fixation, tissues were processed into paraffin via sequential dehydration and perfusion with paraffin under vacuum. Brains were then embedded in paraffin blocks, cut into 6 µm sections and mounted on glass slides for immunofluorescence staining.

### Osmotic Pump Implantation for Intracerebroventricular (ICV) Drug Delivery

For PK and PD studies, osmotic minipumps (ALZET, model 1004) were assembled with the catheter and brain infusion cannula provided in the manufacturer’s brain infusion kit which delivers compounds at 0.11µL/h. Pumps were filled under sterile conditions with either GNE2861 (6 mM; MedChemExpress, HY-12632) or vehicle (DMSO-D2438, Sigma-Aldrich) prepared in 10% (2-hydroxypropyl)-β-cyclodextrin (MedChemExpress, HY-101103). Following filling, pumps were primed by incubation in sterile normal saline in 50 mL conical tubes at 37°C for 48 hours prior to implantation.

For PK analysis, brain infusion cannula connected to the pump was implanted into the right lateral ventricle (coordinates relative to bregma: -1.0 mm anterior–posterior, -0.4 mm medial-lateral,-2.4 mm dorsal-ventral from dura) and secured to the skull using C&B Metabond Quick Adhesive Cement System, Parkell (Cat # S380). The pump was implanted subcutaneously on the back, slightly posterior to the scapulae. At 24 days post-implantation, cerebrospinal fluid (CSF) and hippocampal tissue were collected (n = 3) and analyzed by mass spectrometry to determine absolute compound concentrations.

For PD studies, mice received α-synuclein PFF injections into the medial septum (coordinates relative to bregma: +0.9 mm anterior-posterior, 0.0 mm medial-lateral, -3.85 mm dorsal-ventral from dura). Immediately following PFF injection, osmotic pumps and intracerebroventricular cannula were implanted as described above. Mice were euthanized 24 days post-injection for downstream pathological analyses. Experimental groups consisted of age- and sex-matched cohorts (n = 6 per group) receiving either vehicle or GNE2861. Mice were weighed pre, post-surgery and at the time of endpoint collection.

### GNE2861 LC/MS quantitation

Absolute quantitation of GNE2861 was accomplished using liquid chromatography-mass spectrometry (LC/MS). An external calibration curve was prepared from a high concentration of the drug (GNE2861, HY-12632, MedChemExpress) at 10 µg/mL followed by eight additional points prepared via a half-log serial dilution.

Hippocampi from each mouse were placed in a bead ruptor tube (19-627, Revvity) and weighed. Brain samples and 10 µL of each curve point were extracted in 1 mL of an 80% methanol (A456, Fisher Scientific) (v/v) solution. The solution was spiked with D3 octanoyl-carnitine (HY-139392S, MedChemExpress) such that the final concentration in solution was 5 ng/mL. After solvent addition, each sample was vortexed for 10 s, homogenized for 30 s at 6 m/s at 6°C ± 2°C (cooled via liquid nitrogen), and sonicated for 5 min in a water bath sonicator. Samples were incubated on wet ice for 1 hour, then centrifuged for 10 min at 4°C at 17,000 *xg*. 850 µL of supernatant was transferred to a 1.5 mL tube and the centrifuge step was repeated to fully clear the protein pellet. After the second spin, 800 µL of supernatant was collected to a fresh 1.5 mL tube and dried under vacuum. The dried metabolite extracts were resuspended by adding 40 µL of acetonitrile (A955, Fisher Scientific), followed by 10 s vortex and 5 min water bath sonication. An additional 40 µL of water (W6, Fisher Scientific) containing D8 Tryptophan (DLM-6903, Cambridge Isotope Laboratories) at 1 µg/mL was added to each sample such that the final resuspension solvent composition was 50/50 acetonitrile/water (v/v). After water addition, the 10 s vortex and 5 min water bath sonication was repeated. Samples were then centrifuged for 10 min at 4°C at 17,000 *xg* before transferring 70 µL of supernatant to an autosampler vial with insert (03452247, Thermo Fisher).

Samples and standards were analyzed with an Agilent 6470 triple quadrupole mass spectrometer coupled with an Agilent ultra-high performance liquid chromatography 1290 Infinity II. 2 μL of each sample was injected and separated using a 12-minute gradient on a Cortecs T3 Column (1.6 μm, 2.1mm × 150mm, 186008500, Waters, Eschborn, Germany) combined with a Cortecs T3 VanGuard pre-column (1.6 μm, 2.1 mm × 5 mm, 186008508, Waters). Mobile phase A consisted of LC/MS grade water (W6, Fisher) with 0.1% formic acid (A117, Fisher Scientific). Mobile phase B consisted of 99% Acetonitrile (A955-4, Fisher), 1% LC/MS grade water (W6, Fisher) and 0.1% formic acid. Column temperature was kept at 50°C, flow rate was held at 0.4 mL/min, and the chromatography gradient was as follows: 0-2 min held at 0% B, 2-7.1 min ramp from 0 to 99% B, 7.1-9 min, then held at 99% B. At 9.1 min flow was switched to 100% A at 0.6 mL/min and held until 11 min. By 11.1 min, flow rate was reduced to 0.4 mL/min and allowed to stabilize at the flowrate until 12 min. Gas flow at 13 L/min at 200°C, sheath gas flow at 12 L/min at 325°C and the nebulizer was set to 45 psi. Capillary voltage was +3000 and nozzle voltage was +500. Data was acquired using dynamic multiple reaction monitoring (dMRM) including at least two transitions per compound.

Peak picking and integration were performed using Skyline Software (v 25.1.0.237; RRID:SCR_014080). D3 octanoyl-carnitine was used as an internal standard to assess variability induced by extraction. D8 Tryptophan was used as a measure of run-to-run variability during the course of the LC/MS analysis. The relative standard deviation of each internal standard across all samples was 9.2% and 3.2% respectively, indicating low extraction induced variability and highly stable LC-MS performance.

The resulting peak areas from the GNE2861 calibration curve were used to generate a linear regression for quantitation of the drug in the tissue samples.

GNE2861 concentrations in mouse brain were measured as ng/mg tissue and converted to nM assuming 1 mg of brain tissue ≈ 1 μL, based on standard pharmacokinetic conventions^97^.

### Human Brain Tissue

Substantia nigra paraffin-embedded tissues from 5 idiopathic Lewy body disease brains and 7 healthy matched controls were examined. All procedures were done in accordance with local institutional review board guidelines of the Van Andel Institute Brain Bank, Van Andel Institute, Grand Rapids, MI, USA (RRID:SCR_026035). Written informed consent for autopsy and analysis of tissue sample data was obtained either from patients themselves or their next of kin. Tissues were selected based on neuropathological diagnoses. Non-pathological cases were balanced by age, sex, and PMI.

### Immunofluorescence

Slides were de-paraffinized with 2 sequential 5-minute washes in xylenes, followed by 1-minute washes in a descending series of ethanol: 100%, 100%, 95%, 80%, 70%. Slides were then incubated in deionized water for one minute prior to transfer to the BioGenex EZ-Retriever System where they were incubated in antigen unmasking solution (Vector Laboratories; Cat# H-3300) and microwaved for 15 minutes at 95°C. Slides were allowed to cool for 20 minutes at room temperature and washed in running tap water for 10 minutes. Slides were washed 5 minutes in 0.1 M Tris (diluted from 0.5 M Tris made from Tris base and concentrated hydrochloric acid to pH 7.6), then blocked in 0.1 M Tris/2% fetal bovine serum (FBS) for 1 hour. Slides were incubated in primary antibody in 0.1 M Tris/2% FBS in a humidified chamber overnight at 4°C; pS129 α-synuclein (EP1536Y, Abcam ab51253, RRID: AB_869973, 1:5000); pS129 α-synuclein (81A), BioLegend 825701, RRID: AB_256489, 1:1000; Guinea Pig NeuN (Synaptic Systems 266 004, RRID: AB_2619988, 1:1000), pPAK(GII), Cell Signaling 3241, RRID: AB_2158623, 1:500). Primary antibody was rinsed off with 0.1 M tris for 5 minutes and incubated with secondary antibodies (Goat anti-Rabbit IgG 647, Thermo A-21244, RRID: AB_2535812, 1:1000); (Goat anti-Guinea Pig IgG 568, Thermo A-11075, RRID: AB_2534119, 1:1000) on slides in the dark for 3 hours at room temperature, rinsed and washed 0.1 M Tris for 10 minutes three times, then mounted with coverglass in ProLong gold with DAPI (Invitrogen, Cat#P36931). Fluorescent slides were imaged at 20x magnification on a Zeiss AxioScan 7 microscope or on the ImageXpress Confocal HT.ai High-Content Imaging System at 60x magnification.

### Mouse Brain Segmentation and Registration

#### Segmentation

Stained slides were scanned on a Zeiss AxioScan 7 at 20x magnification and imported into QuPath v0.5.0 (RRID: SCR_018257)^98^ for analysis. A pixel classifier thresholding fluorescent intensity for the pS129 α-synuclein channel was applied to each section, picking up positive pathology signal for quantification. Signal intensity was optimized by adjusting the min/max display settings. A threshold with similar settings was applied across cohorts. Individual cells were identified using the cell detection feature in QuPath that detects cells based on DAPI. Cell detection parameters such as background radius, sigma, and threshold were adjusted to optimize cell detection across brain sections. A cell expansion parameter was set to 5 µm to include the full nuclear and perinuclear NeuN staining and establish cell bounds that capture all pathology within the soma of cells. An object classifier identifying cells positive for pS129 α-synuclein was applied to sections. QuPath was trained on a subset of annotations to ensure proper classification of positive cells based on signal intensity. Once sufficiently accurate, the classifier was loaded onto the entire brain image to classify all detected cells. The subcellular spot detection feature was used to identify pS129 α-synuclein signal within positive pS129 α-synuclein cells for the inclusion area measure. The same fluorescent intensity level set for the pixel thresholder was applied for subcellular spot detections.

#### Mouse Coronal Section Brain Registration

Images were registered to the Allen Brain Atlas CCFv3 using a modified version of the QUINT workflow (RRID: SCR_023856)^41^. An RGB image of each section was exported from QuPath as a PNG, down sampled by a factor of 12, to use for spatial registration. Segmentation was created by exporting a color-coded image of classified pixels or cells on a white background for use as the segmentation input in Nutil (RRID:SCR_017183)^99^. Brain images were uploaded to the web version of DeepSlice (RRID: SCR_023854)^100^ and the deep neural network was used to automatically align sections to regions of the atlas. Preliminary alignment was exported from DeepSlice as an XML that was then uploaded to QuickNII (RRID: SCR_016854)^101^ for further refinement. Following the spatial registration of the mouse brain sections to the Allen Mouse Brain Atlas CCFv3 in QuickNII, a JSON file was saved for use in VisuAlign (RRID:SCR_017978). Brain sections were imported into VisuAlign to fine tune the registrations to match regions of interest. Anchor points were generated in the atlas overlay and moved to the corresponding location on the brain section via non-linear transformations. Markers were placed around the contour of the brain section first with markers refining the inner structures applied second. Final alignments were exported as FLAT and PNG files for use in Nutil^99^.

Nutil (RRID:SCR_017183)^99^ was used for the quantification and spatial analysis of the identified cell types in specific regions of the mouse brain. Each segmentation (pixels and cells) was run through Nutil by uploading the input files generated from the masks pulled out in QuPath, a JSON anchoring file from QuickNII, and a folder with the FLAT and PNG files from VisuAlign containing the transformed registration. Individual classes were identified for quantification via their HTML color code assigned in QuPath. Nutil generated object area, region area, area occupied, and object counts from each individual classification within each region of the ABA CCFv3 using the registration from QuickNII and VisuAlign. The outputs are used in modeling and in N2U to generate anatomical heatmaps. The render of registered sections was generated in Meshview (RRID:SCR_017222)^101^.

#### 3D Mouse Brain Registration and Segmentation

The α-synuclein channel from the SPIM data set (tiff image stack) was imported into Fiji/ImageJ v1.54f^102^ for preprocessing (manual threshold and masking with CLIJ2) and saved as a multipage tiff prior to registration to the ABA CCFv3 using BrainGlobe’s BrainReg software^103^. The following key parameters were used for registration: voxel size: 4.0 z, 1.8 xy; orientation: sal; geometry: full brain; atlas: 10 µm; save original orientation: True. The exported brain atlas label values were remapped to 16-bit space for easier downstream analysis using a custom python script and then up sampled in Fiji to half the size of the original dataset in xyz. The hemisphere mask generated by BrainReg was transposed in yz with CLIJ2^104^ and used to uniquely label all brain regions. The original SPIM dataset in the pS129 α-synuclein channel was relabeled using a 16-bit atlas, and the transposed hemisphere images were all imported into Imaris (Oxford Instruments) as a single ims file. The relabeled 16-bit atlas was additionally imported as a label/segmentation image for automatic conversion into Surface objects. The pS129 α-synuclein signal was then segmented with the Surfaces module using automatic filtering parameters and a surface grain size of 0.5 µm. Overlap between brain region surfaces and segmented α-synuclein surfaces was calculated with Imaris’ Object-Object statistics option. Quantitative information was exported for further analysis.

### Anatomical Analysis Heatmaps

#### Nutil-to-Usable (N2U)

Nutil-to-Usable is an R-based shiny app developed by bioinformaticians at Van Andel Institute designed to aid the plotting of regional data to anatomical heatmaps based on the Allen Brain Atlas (RRID:SCR_024753). Outputs from Nutil were uploaded with data from the left and right hemispheres submitted separately. An annotation file is generated allowing for labelling of metadata for each section’s associated Nutil file (mouse id, sex, treatment, post-injection interval, etc.). Checkpoint files compiling the raw Nutil files and metadata were downloaded to facilitate faster upload times when working downstream in the app. Settings were then defined to determine the level of summary and variable of interest sought to average. ‘Load’, defined as the area of pathology/area of the region, was set as variable of interest when plotting for total and cell body inclusion area occupied. To plot neurite area occupied, checkpoint files from total and inclusion were combined and uploaded with the app variable of interest set to ‘Neurite Load’. The app runs calculations to subtract cell body inclusion area from total area for each region and run relevant calculations to divide by region area resulting in neurite area occupied. Level of resolution for the atlas is set to ‘daughter’ (more specific, individual layers and subregions shown), or ‘parent’ (more general, layers compiled into single region). Grid Heatmaps and plotting matrices are developed in this step.

#### Mouse Brain Heatmaps (MBH)

Plotting matrices were uploaded to MBH to generate anatomical heatmaps. MBH plots the compiled data for the parameter of interest represented in the plotting matrix set by N2U. Applicable coronal section figures were listed in the app by number based on the ordering of Allen Brain Atlas sections. Regional Resolution for the plots was set to either daughter (by layer) or parent (compiling layers). Plots were downloaded as svgs.

#### Statistical Analysis

For analyses assessing differences in percent area occupied between timepoints, data were stratified based on brain hemisphere (ipsilateral/contralateral), brain region, and MPI. Regions with zero variance across all timepoints were filtered out. To maintain flexibility and avoid reliance on strong parametric assumptions, robust linear regressions via the MASS package, with the ranked percent area occupied as the outcome and MPI as the explanatory variable, were used (https://cran.r-project.org/web/packages/MASS/index.html, RRID: SCR_019125). All models were adjusted for sex and daughter regions when the number of daughter regions was 2 or greater. Effect sizes, including confidence intervals were estimated using the R package emmeans (R package version 1.8.3, URL: https://CRAN.R-project.org/package=emmeans, RRID: SCR_018734). Significance was determined using second-generation P values based on a null interval of ± 5% difference with 95% confidence intervals. Only second-generation P values equal to 0 were considered significant.

#### Quantification of Brain Pathology Following Alzet Pump-Mediated Compound Delivery

Brain sections corresponding to hippocampal levels (Allen Brain Atlas CCFv3 sections 72, 82, and 88) were collected and stained for pS129 α-synuclein and NeuN. Regions of interest (ROIs) corresponding to field CA1, field CA3, and the dentate gyrus (DG) were annotated. A pixel classifier threshold for pS129 intensity was applied to identify the total pS129-positive area within each ROI.

NeuN-positive cells were identified using the Cell Detection tool in QuPath, with segmentation parameters adjusted to achieve consistent detection across brain regions. An object classifier was then trained and applied to identify positive cells for pS129 α-synuclein pathology.

Subcellular detection was used to identify pS129 α-synuclein-positive pixels within classified cells, which were defined as the pS129 α-synuclein cell body area. Neuritic pathology was calculated by subtracting the cell body area from the total pS129 α-synuclein-positive area within each ROI. The percentage area occupied by pathology within each hippocampal subfield was then calculated and plotted.

For quantification of pPAK intensity, brain sections adjacent to those used for pathology quantification, corresponding to hippocampal level 82 of the Allen Brain Atlas CCFv3 were used for quantification of phosphorylated Group II PAK (pPAK) signal. Sections were stained for pPAK and NeuN, and regions of interest (ROIs) corresponding to field CA1, field CA3, and the dentate gyrus (DG) were annotated. NeuN-positive cells were identified using the Cell Detection tool in QuPath, and the mean pPAK intensity within NeuN-positive cells was quantified for each ROI. Background fluorescence was measured from an area within the corresponding ROI that was devoid of neuronal cell bodies and was subtracted from the mean pPAK intensity for each section. Background-corrected pPAK intensities were grouped according to treatment (vehicle or GNE-2861) and expressed as fold-change relative to the vehicle group.

### Mouse Brain Microdissection and Cerebrospinal Fluid Collection

Cerebrospinal fluid (CSF) was collected from deeply anesthetized mice by carefully removing the skin and muscle layers around the cisterna magna. Once the cisterna magna was exposed, the area was wiped with a cotton swab to remove any blood. A needle was used to puncture a hole through the membrane of the cisterna magna and CSF was collected using a gel loading tip. The CSF was spun down to remove any possible contaminants. Next, the brains were removed from the skull and placed in a shallow petri dish with PBS on ice. The brains were cut into 2 mm coronal sections and hippocampi from entire rostrocaudal axis were dissected and flash frozen for downstream analysis.

### GeoMx Spatial Transcriptomics

#### Tissue Preparation

Spatial transcriptomics was performed using the nanoString GeoMx® Digital Spatial Profiler. Sections were cut at 6 μm thickness and mounted on plus-charged slides (Epredia Colormark Plus CM-4951WPLUS-001). Slides were baked at 60°C for 1 hour and stored at 4°C in a vacuum sealed container containing desiccant for up to two weeks. All subsequent steps were performed using RNase-free conditions and DEPC treated water. Slides were de-paraffinized with 3 sequential 5-minute washes in xylenes, followed by 2 washes in 100% ethanol for 5 minutes, 1 wash in 95% ethanol, and 1 wash in 1x PBS. Target retrieval was performed in target retrieval reagent (10x Invitrogen 00-4956-58 EDTA pH 9.0) diluted to 1x in the BioGenex EZ-Retriever System for 10 minutes at 95°C. Slides were then washed with 1x PBS for 5 minutes. Slides were then incubated in 0.1μg/mL proteinase K (Invitrogen 25530-049) for 10 minutes at 37°C and washed in 1x PBS for 5 minutes at room temperature. Slides were post fixed for 5 minutes in 10% neutral buffered formalin followed by two washes in NBF stop buffer (24.5g Tris base and 15g Glycine in 2L DEPC water) for 5 minutes each and one wash in 1x PBS for 5 minutes. Slides were then incubated with hybridization probes (nanoString Cat# 121401103) diluted in Buffer R (provided in the GeoMx RNA Slide Prep FFPE-PCLN kit, catalog # 121300313) in a hybridization oven at 37°C for 16-20 hours.

Following probe incubation, slides were washed with stringent washes (equal parts formamide and 4x SSC buffer) at 37°C twice for 25 minutes each. Then slides were washed twice in 2x SSC buffer. Slides were incubated in 200 μL buffer W (provided in the GeoMx RNA Slide Prep FFPE-PCLN kit, catalog # 121300313) for 30 minutes and incubated in morphology markers (GFAP-488, ThermoFisher Scientific 53-9892-82, RRID:AB_10598515, 1:400; HuC/HuD (16A11) ThermoFisher Scientific A21271, RRID:AB_221448,1:500; NeuN, Millipore ABN78, RRID:AB_10807945, 1:1000) at 4°C overnight. The first four mice were characterized with NeuN only as the morphology marker. NeuN did not show efficient staining of several subcortical regions, so HuC/HuD were optimized for these regions and the last two mice included NeuN and HuC/HuD antibodies to enable segmentation of subcortical neuron types. Slides were washed 4 times in 2x SSC buffer for 3 minutes each wash. Slides were then incubated with secondary antibodies (GαRb 647, ThermoFisher Scientific A21244, RRID:AB_2535812, 1:1000; GαIgG2b 647 Thermo A21242, RRID:AB_2535811, 1:1000) and nuclei marker Syto83 (Thermo Scientific S11364, 1:1000) in Buffer W for 1 hour at room temperature in a humidified chamber. Slides were washed 4 times in 2x SSC buffer for 3 minutes each and placed in the nanoString GeoMx® DSP instrument.

Syto83 immunofluorescence was utilized for autofocus of GeoMx imaging. Immunofluorescence for GFAP was used in identification of morphological markers to aid in fitting to the Allen Brain Institute’s mouse brain atlas. NeuN and HuC/HuD were used for segmentation. Prior to region-of-interest (ROI) generation, the slide image was exported, and individual brains were registered to the Allen Brain Atlas CCFv3^105^ using the mouse brain registration protocol. Images of registered brains were imported onto the DSP instrument and fit exactly to the slide image to enable accurate anatomical selection of ROIs. ROIs were generated using the polygon or circle tool which aligned with the correct brain region. Each ROI was segmented into one area-of-illumination (AOI) to detect neurons within the designated ROIs. Probe identities in each segment were captured via UV illumination and moved to a 96-well plate.

### NGS Library Prep and Sequencing

Library preps and sequencing were performed by the Van Andel Genomics Core. Illumina Novaseq 6000 was used for sequencing with a read length of 27 for both reads with reverse sequence orientation in the readout group plate information. Plates were dried down and rehydrated in 10 µL nuclease-free water, mixed and incubated at room temperature for 10 minutes. PCR was performed on samples as described in the nanoString GeoMx® DSP Readout User Manual using 2 µL PCR master mix, 4 µL primer from the correct wells, and 4 µL resuspended DSP aspirate. KAPA beads (KAPA Pure beads, Roche Cat# 07983298001) were warmed to room temperature for 30 minutes. Libraries were pooled and KAPA beads were added to each pool at a 1.2X ratio to the final pool volume. Two KAPA bead clean ups were performed and pooled libraries were eluted in 24 µL elution buffer. Negative and positive control pools were eluted into 10 µL elution buffer. Quantity of the pools were assessed using the QuantiFluor® dsDNA System (Promega Corp.). Pools were diluted to 5 ng/µL and quality and size are assessed using Agilent DNA High Sensitivity chip (Agilent Technologies, Inc.) on the Bioanalyzer. Sequencing was performed at 100 reads/µm^2^. Paired end 50 base paired sequencing. Base calling was done by Illumina RTA3 and output was demultiplexed and converted to fastq format with bcl2fastq v1.9.0. ( RRID:SCR_015058). Fastq files were then converted to Digital count conversion files (DCC) using GeoMx NGS Pipeline v2.3.3.10. Sequencing reads were trimmed to 27 bp to reflect GeoMx probe length.

### Quality Control and Data Analysis

In the PANGEA experiment, 717 segments were captured in total from 7 mice (3 male and 4 female; age=3 months); 16 segments failed quality control (QC) analysis and were removed. One segment was removed for trimmed reads < 80%, stitched reads < 80%, and aligned reads < 75%. 14 additional segments were removed for sequence saturation below 50%. One additional segment was removed for < 3% of genes being detected above the limit of quantification (LOQ). Of the 20,175 gene targets contained in the GeoMx Mouse Whole Transcriptome Atlas, 19781 were included in the downstream analyses following QC analysis. 1 gene target was removed as a global outlier (Grubbs test, P<0.01), and 10 were removed as local outliers (Grubbs test, P<0.01). Of the remaining gene targets, 19,781 gene targets were detected above the LOQ in at least 1% of segments and therefore were selected for further analysis. An additional 6 gene targets were identified as interesting *a priori* and were retained in the study for additional analysis, regardless of QC performance. Sample size was determined based on pilot data. No statistical method was used to determine sample size. Experiments did not involve multiple experimental conditions.

The PANGEA app was developed as a server-free web application designed to visualize and analyze brain region gene expression data. Developed using the Lume Core platform by Lume VR Limited (https://www.lumevr.com, Oxford, UK; contact), PANGEA offers a novel, ultra-low-maintenance solution with no ongoing operational costs, ensuring long-term accessibility and sustainability. This server-free architecture means that all data uploaded by end-users remains solely on their local machines, never shared or stored externally, which allows for secure analysis of proprietary or sensitive data. The platform hosts spatial gene expression data from mouse neuroanatomical regions, mapped onto the ABA CCFv3, enabling researchers to explore transcriptomic profiles in anatomically defined brain regions with ease and security.

In the mouse GeoMx experiment data used for Fig. 6H and 8G, and the human GeoMx experiment used for Fig. 8G, post-QC data was downloaded from the publicly available data previously published by our group and utilized without additional alterations (https://doi.org/10.5281/zenodo.10729767)^40^. This data was originally produced using similar QC analyses as described above. For detailed QC parameters of this data see https://github.com/Goralsth/Spatial-transcriptomics-reveals-molecular-dysfunction-associated-with-cortical-Lewy-pathology (DOI: 10.5281/zenodo.10732492).

### Primary Neuron Culture from Wildtype Mice

Primary neuron cultures were prepared from embryonic day 18 (E18) CD1 (strain 022; RRID: IMSR_CRL:022) mice. Brains were gently removed from the embryos and placed into a petri dish filled with ice-cold, sterile Hibernate Medium (Cat#. A1247601, Gibco) The hemispheres were gently separated, and the meninges, thalamus, striatum, brainstem, and hippocampus were removed. Hippocampi were isolated, pooled and digested in papain solution (20 U/mL Cat# LS003126, Worthington) and then treated with DNase I (Cat# LS006563, Worthington) to remove residual DNA. The tissue was then washed with pre-warmed Neurobasal media (Cat# 21103049, Gibco), mechanically dissociated, and strained through a 40 μm cell strainer. The cell suspension was pelleted at 1000 x *g* for 5 minutes, resuspended in 2 mL of neuron media (Neurobasal media containing 1% B27, 2 mM GlutaMAX, and penicillin-streptomycin), and gently mixed. The dissociated neurons were seeded on poly-D-lysine (Cat# P0899, Sigma) coated 96-well culture plates (Cat# 655090, Greiner) at 17,000 cells/well. Cells were maintained at 37°C at 5% CO_2_.

### Primary neuron culture from *Pak5/6* knockout mice

Primary neuron cultures were prepared from postnatal P (0-1) B6;129-*Pak6^tm1Amin^ Pak5^tm1Amin^*/J mice (Strain #:015825; RRID:IMSR_JAX:015825). Heterozygous mice were bred, and hippocampi from individual pups were processed independently to generate primary neuron cultures, with each pup treated as a separate biological replicate. Each hippocampus was cultured as described in the primary neuron culture from wildtype mice. Media suitable for postnatal culture was used; Hibernate media-A (Cat# A1247501, Gibco) and Neurobasal-A (Cat# 10888022, Gibco). The tail from the pups were used to extract genomic DNA and confirm the genotype of the pups using the following primers, *Pak5* WT forward, 5’-GCTTCCTCAGATCCATCCAAGGT-3’; *Pak5* mutant, 5’-CTTCCTGACTAGGGGAGGAGT-3’; Pak5 reverse, 5’-AGATGCATTGAGTGCTGGGGAA-3’; Pak6 WT forward, 5’-TCAGTTATCAGCTCCAACACCCTG -3’; *Pak6* mutant forward, 5’-GCTACCGGTGGATGTGGAATGTGT-3’; *Pak6* reverse, 5’-GAGGAAACCCCAGGTCATATACCT-3’.

### Kinase Inhibitor Screen

All 12 compounds for the kinase screen and 7 PAK inhibitors were procured from MedChemExpress (listed in the Key Resources Table). Each compound was reconstituted in dimethyl sulfoxide (DMSO-D2438, Sigma-Aldrich) to create a stock solution at its maximum solubility . The three treatment doses were determined based on the compounds’ IC50 values for the target kinase. The stock solutions were further diluted with DMSO and aliquoted in a 96-well plate to the desired concentration for the treatment ensuring the final concentration of DMSO remains <0.1% at all tested dosages. On the day of treatment, the diluted compounds were mixed with neuron media and added to the primary neurons cultured in 96-well plates.

### Immunocytochemistry

Neurons were fixed with prewarmed 4% paraformaldehyde/4% sucrose in phosphate buffer saline (PBS) for 15 mins, followed by 5 washes with PBS. Cells were then permeabilized with 0.3% Triton X-100 in 3% bovine serum albumin (BSA) for 15 mins followed by a 3X wash with PBS. For the detection of detergent-insoluble α-synuclein, neurons were fixed with paraformaldehyde containing 1% Triton X-100 to extract soluble proteins. Cells were blocked with 3% BSA for 1 hour before incubation with primary antibodies (MAP2, Abcam AB5543, RRID: AB_571049, 1:5000; NeuN, Millipore MAB377, RRID: AB_2298767, 1:1500; pS129 α-synuclein (81A), BioLegend 825701, RRID: AB_256489, 1:2000; pPAK(GII), Cell Signaling 3241, RRID: AB_2158623, 1:500) at room temperature for 2 hours. Cells were washed 5X with PBS and were then incubated with fluorescent secondary antibodies (Goat anti-chicken-488, Thermo Scientific-A11039, RRID: AB_2534096; Goat anti mouse IgG2a-546, Thermo Scientific-A21133, RRID: AB_2535772; Goat anti mouse IgG1-647, Thermo Scientific-A21240, RRID: AB_2535809) for 1 hour at room temperature in dark, followed by 5 washes with PBS. All fluorescent secondary antibodies were used at a dilution of 1:500. Cells were then stained, incubated with DAPI, 1:10,000 in PBS. The plates were sealed and stored at 4°C until imaging.

### Imaging and Image Analysis

96-well plates of stained primary neurons from the kinase screen were imaged on ImageXpress Confocal HT.ai High-Content Imaging System at 10X magnification. For pPAK analysis of primary neurons derived from PAK5/6 KO mice, plates were imaged at 10X magnification on the ImageXpress Confocal HCS system. 9 fields-of-view per well were imaged and imported into MetaXpress Analysis software (V3.7). MAP2/pS129 α-synuclein positive signal was segmented using ‘Find Blobs’ tool with minimum width of 1 µm and maximum widths of 30 µm and 50 µm for MAP2 and pS129 α-synuclein, respectively. NeuN positive nuclei were segmented with minimum width of 9 µm and maximum width of 45 µm. The threshold for each channel was kept the same for the entire plate and adjusted to only quantify true positive signal to background noise. Total area covered for MAP2 and pS129 α-synuclein and number of NeuN positive nuclei was quantified for each image. 96-well plates from the suite of PAK inhibitor screen were imaged on Zeiss Cell Discoverer 7 at 20X magnification with 0.5X digital zoom. 10 ROI per well were imaged. Images were analyzed on Zeiss Blue Analysis software (V3.7). The ‘Automatic segmentation’ tool was used to segment each signal. For MAP2 and pS129 α-synuclein, a ‘Minimum Object Size’ of 1 was chosen and the threshold was set for each plate manually to segment most of the MAP2 and pS129 α-synuclein signal. For NeuN and DAPI, a ‘Minimum Object Size’ of 200 was chosen and minimum and maximum thresholds were set manually for each plate to segment the NeuN and DAPI signal. Background thresholds were adjusted for true positive to background signal. NeuN and DAPI positive cells were counted, and the areas of MAP2 and pS129 α-synuclein were measured for each image. Data from all images/wells were pooled, averaged, and normalized to control wells and represented as fold change.

For pPAK puncta analysis in mouse tissue and primary neuron cultures, slides and 96-well plates were imaged using the ImageXpress (IX) Confocal HT.ai High-Content Imaging System at 60x magnification. Images were quantified on the MetaXpress (MX) Analysis software. NeuN positive nuclei for mouse amygdala, NeuN/HuC/HuD for mouse substantia nigra (SN) and DAPI for primary neuron culture and were identified using the ‘Find Blob’ feature by training the MX for size and fluorescence intensity of the signal over background. For primary neurons, MAP2-positive DAPI was segmented as neuronal nuclei as an additional step. The cell body was segmented by subtracting DAPI from MAP2 in primary neurons and NeuN in the mouse amygdala and NeuN/HuC/HuD in mouse SN. pS129 α-synuclein was segmented using the ‘Auto Threshold’ module and trained to identify true signal over background. pPAK puncta were segmented with the ‘Find Blob’ feature by training for size and fluorescence intensity of the true signal over background. pPAK puncta integrated intensity and number of puncta, and pS129 α-synuclein total area for each neuronal cell body were quantified. Cells were classified as positive and negative for pS129 α-synuclein stain, and the percentage of neurons with pPAK puncta in pS129 α-synuclein positive and negative cells was calculated.

### Statistical Analysis

GraphPad Prism software version 10.2.2 (GraphPad Software Inc., La Jolla, CA, USA) was used for statistical analysis for primary neuron culture experiments.

### Computational Modeling of α-Synuclein Pathology Spread

#### Network spreading dynamics

Our model assumes that α-synuclein pathology proceeds bidirectionally in a linear fashion along the brain’s structural connectome. The structural connectivity matrix W was derived from the weighted region-to-region connectome generated by Knox et al.^106^, which aggregates viral tracer experiments from the Allen Mouse Connectivity Atlas into a directed structural connectivity matrix over Allen CCFv3 brain regions. Thus, W*_ij_* represents the strength of the projection from region *j* to region *i*. From this weighted and directed connectivity matrix, we constructed the out-degree graph Laplacian *L* = *D* − W, where *D* is the diagonal out-degree matrix.

Pathology spread was modeled as a continuous-time linear diffusion process on the structural connectome,

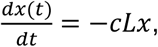

where *x*(*t*) denotes the regional pathology at time *t*, *c* is the diffusion rate constant, and *L* is the graph Laplacian. The closed-form solution of this linear differential equation is given by the matrix exponential. The predicted pathology arising from anterograde and retrograde spread was therefore obtained as

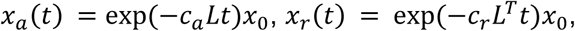

where *c_a_* and *c_r_* are diffusion rate constants for spread in the anterograde and retrograde directions, respectively, *L* represents the out-degree graph Laplacian, *L^T^*denotes its transpose,

*t* is time, and *exp*(×) denotes the matrix exponential, which is the closed-form solution of the linear diffusion equation. The vector *x*_0_ represents initial pathology and contains a 1 in the ipsilateral caudoputamen and zeros elsewhere.

To compare model predictions with experimentally measured pathology, predicted regional pathology values were log_10_-transformed before linear regression, as both measured and predicted pathology spanned several orders of magnitude. The bidirectional network diffusion model therefore took the form

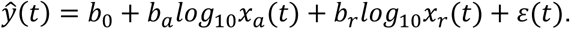

Here, *b*_0_ is the intercept, *b_a_* and *b_r_* represent the relative contributions of anterograde and retrograde spread, respectively, and *ε*(*t*) is the residual error. We used the ‘optim’ function in R to identify the values of *c_a_* and *c_r_* that maximized the average Pearson correlation coefficient between actual and predicted pathology over the time points *t* = 0.1, 0.2, 0.3, 0.5, 1, 3, 6 and 9 MPI.

### Network Model Validation

We used two established procedures to assess the performance of our predictive model^29,31,33^. First, to evaluate the model’s specificity to the experimental seed site, we randomly selected 500 alternate seed regions and redefined the vector *x*_0_ accordingly. We then refit the model for each alternate seed region to obtain a distribution of Pearson’s r values. For each time point, we obtained a one-tailed, nonparametric *p* value that reflects the proportion of times that the fit obtained from the actual seed site was stronger than that resulting from fitting the model to alternate seeds.

The linear network diffusion model used here assumes that the spread of pathological α-synuclein proceeds bidirectionally. Model parameters, including diffusion rate constants and time-dependent weights for anterograde and retrograde spread, are fit using the available data from all mice at every time point. To statistically compare the out-of-sample performance of our bidirectional network diffusion model to models based on anterograde spread alone, retrograde spread alone, or Euclidean distance, we randomly sampled the data to obtain *y_train_*(*t*) and *y_test_* (*t*) for each time point. Parameters were estimated using the model fitting process described in the preceding section on *y_train_*(*t*), and the model’s performance was evaluated by its fit with *y_test_* (*t*). This procedure was repeated 500 times for each model type to generate a distribution of model fits. Pairwise two-tailed nonparametric tests were used to statistically compare the strength of correlations between actual and predicted pathology obtained by each model type.

### Regional Vulnerability

In our linear network diffusion model, *ε*(*t*) represents residual variance that is not explained by spread along the brain’s structural connectome. Positive values of *ε*(*t*) indicate that a region was occupied with higher levels of pathology than would be predicted by structural connectivity alone, and negative values of *ε*(*t*) indicate that a region accumulated lower levels of pathology than would be predicted by structural connectivity alone. To obtain a measure of regional vulnerability to pathology, we averaged the residual variance across both hemispheres and across the 1, 3, 6 and 9 MPI time points. Earlier time points (0.1, 0.2, 0.3, and 0.5 MPI) were excluded from the measure because model performance was weaker at those time points and patterns of residuals were dissimilar from later time points (**Fig. S6C**).

We evaluated the relationship between regional vulnerability and gene expression to identify genes that may contribute to regional susceptibility to pathology. First, a scaled sigmoidal transformation was used to normalize levels of gene expression across the brain for each gene, such that all expression levels fall on a scale of 0 to 1. A rank inverse normalization transformation was then applied to both regional vulnerability and gene expression data. To identify genes that are associated with regional vulnerability, we computed Pearson’s correlations between regional vulnerability and gene expression for each gene in the PANGEA dataset. p-values were FDR corrected for multiple comparisons.

## ACKNOWLEDGEMENTS

We thank the Van Andel Institute Bioinformatics and Biostatistics Core (RRID:SCR_024762), especially Zachary Madaj, for their assistance with statistical analysis, the Pathology and Biorepository Core (RRID:SCR_022912) for their assistance with tissue sectioning and staining, the Genomics Core (RRID:SCR_022913) for their assistance with GeoMx spatial transcriptomics, the Optical Imaging Core (RRID:SCR_021968), especially Kristin Gallik, for their assistance with imaging and image analysis, the Van Andel Institute Mass Spectrometry Core (RRID:SCR_024903) for PK analysis, the West Michigan Neurodegenerative Diseases (MiND) Program iPSC and Screening Platform (RRID:SCR_027675), and the Vivarium (RRID:SCR_023211). We would like to thank the patients and families who donated tissue to research, without whom this study would not have been possible. Human brain tissue was provided by the Van Andel Institute Brain Bank (RRID:SCR_026035), which is supported by the West Michigan Neurodegenerative Disease (MiND) Program. We acknowledge Lume VR Limited (https://www.lumevr.com, Oxford, UK; contact) for development of the PANGEA app.

This research was funded in part by Aligning Science Across Parkinson’s (ASAP-020616 to M.X.H. and D.S.B.) through the Michael J. Fox Foundation for Parkinson’s Research (MJFF), NIH grant R21-NS116255 to M.X.H and a West Michigan Neurodegenerative Disease (MiND) pathway-to-independence award to N.V (MiP2i.002.NV). Several images were created with BioRender.com. For the purpose of open access, the author has applied a CC BY public copyright license to all Author Accepted Manuscripts (AAM) arising from this submission.

## AUTHOR CONTRIBUTIONS

N.V. and M.X.H. conceived the project. N.V., J.K.B., T.M.G., K.K., L.M., L. Breton, and M.X.H. designed experiments. N.V., K.K., L.M., L. Breton., L.T., J.K.B., C.I., and M.X.H. performed experiments. N.V., J.K.B., T.M.G., K.K., L.M., L. Breton., D.D., L. Brasseur., K.L.B., C.I., R.S.D., D.S.B., and M.X.H. developed experimental protocols, tools, and reagents or analyzed data. N.V., J.K.B, T.M.G., K.K., D.S.B., and M.X.H. wrote the manuscript. All authors reviewed and approved the manuscript.

## DATA AND MATERIALS AVAILABILITY

The data that support the findings of this study together with the code used for data analysis are available here: https://github.com/Goralsth/Synuclein_Progression_Github.

Primary code used to analyze data and generate linear network diffusion models is available here: https://github.com/jkbrynildsen/aSyn_spread.

Immunofluorescence imaging dataset supporting this manuscript are available here: https://doi.brainimagelibrary.org/doi/10.35077/g.1178

Initial conversion of Nutil files to matrix output and grid heatmap generation:

https://github.com/DaniellaDeWeerd/NutilToUsable (RRID:SCR_024753). Brain heatmaps and differential analysis of area occupied: https://github.com/vari-bbc/Mouse_Brain_Heatmap.

The cleaned tabular dataset used to generate the figures and findings of this manuscript is publicly available in the Zenodo repository at https://doi.org/10.5281/zenodo.20648167.

All the resources used are listed in a Key Resource Table publicly available in the Zenodo repository https://doi.org/10.5281/zenodo.22681170.

All lab protocols used in this paper are publicly available on protocols.io:

α-Synuclein Protein Preparation dx.doi.org/10.17504/protocols.io.j8nlkop65v5r/v1

Embryonic/Postnatal Mouse Neuron Culture dx.doi.org/10.17504/protocols.io.14egn68jql5d/v1

Immunocytochemistry dx.doi.org/10.17504/protocols.io.5jyl82ym8l2w/v1

Immunofluorescence Protocol dx.doi.org/10.17504/protocols.io.4r3l22y6jl1y/v2

Stereotaxic Surgery dx.doi.org/10.17504/protocols.io.yxmvm3zy5l3p/v1

Transcardial Perfusion in Mouse dx.doi.org/10.17504/protocols.io.dm6gpbrw1lzp/v1

QUINT Workflow for Fluorescence dx.doi.org/10.17504/protocols.io.4r3l22y6jl1y/v2

LC-MS quantitation of GNE2861 dx.doi.org/10.17504/protocols.io.kqdg3mjpel25/v1

Alzet Pump Placement and Stereotaxic Injections dx.doi.org/10.17504/protocols.io.kqdg3mod1l25/v1

Cerebrospinal Fluid Collection (Terminal) dx.doi.org/10.17504/protocols.io.6qpvr1eybgmk/v1

