## Supplemental Figures for "Molecular analysis of network vulnerability to α-synuclein pathology reveals PAKs as therapeutic targets for Parkinson’s disease"

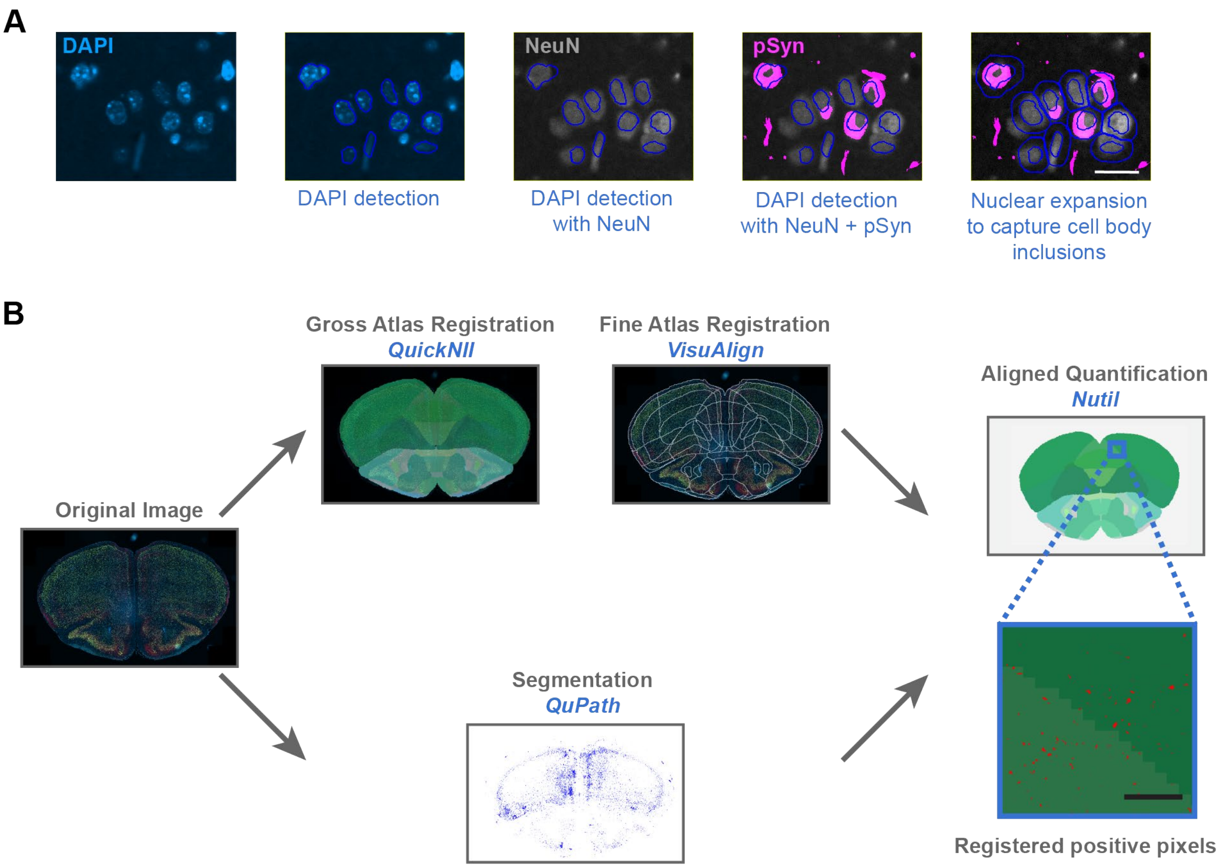


**Figure S1. Registration and segmentation strategy** (A) Demonstration of DAPI based segmentation with 5um cell expansion to capture pathology contained in perinuclear NeuN staining. DAPI in blue, NeuN in grey, and pSyn in magenta. Scale bar = 20 µm. (B) Schematic of the modified QUINT workflow used to analyze brain sections. Following staining, brain sections were digitized. Those digital images were registered to the ABA CCFv3 with QuickNII and VisuAlign software. Segmentation for total, cell body, and neuritic pathology was performed in QuPath. Segmentation and registration were then integrated to generate pathology measures in each anatomical region.

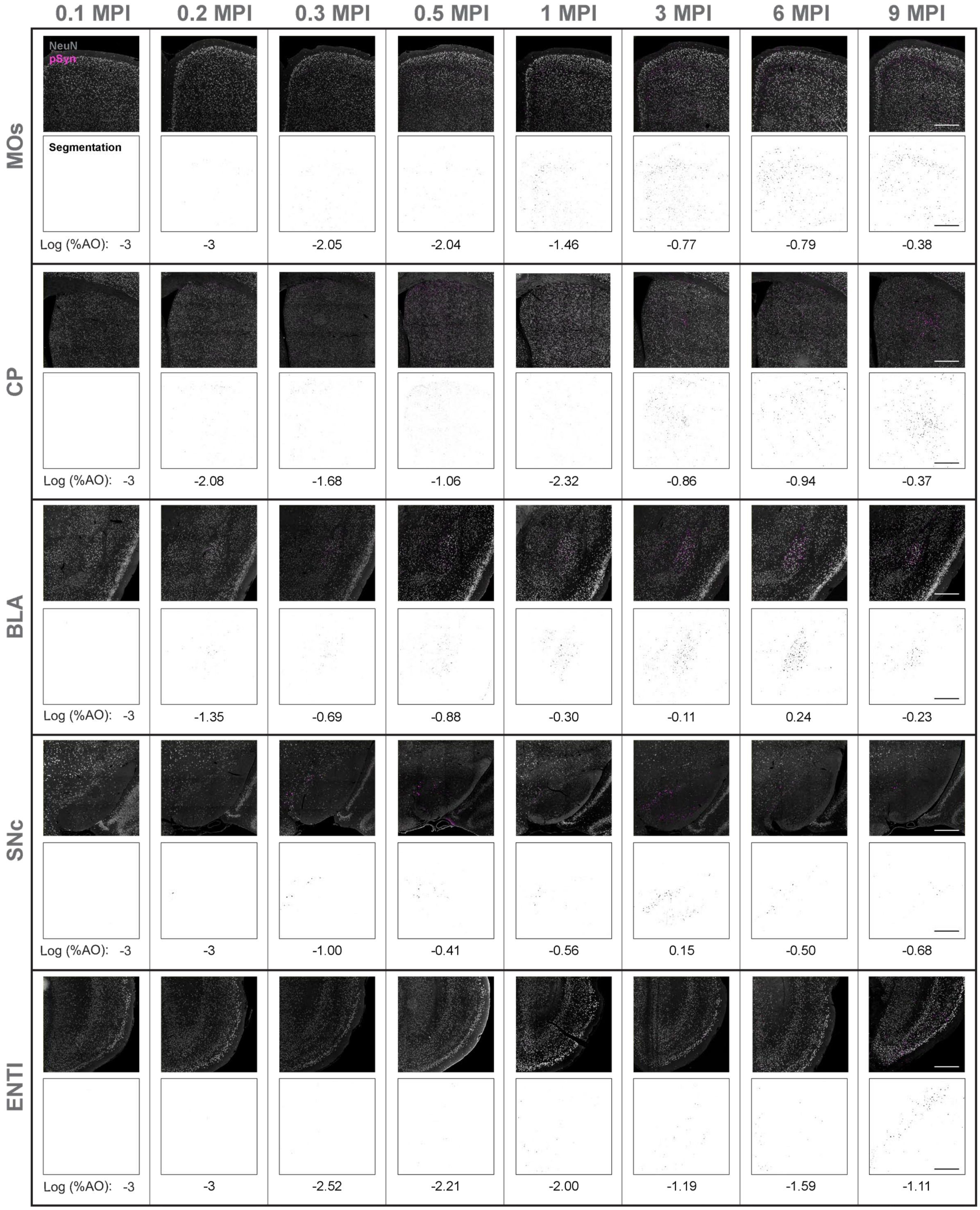


**Figure S2. Regional representative images.** Related to Fig. 1d, zoomed out images of NeuN/pSyn staining are shown for each region across timepoints. The pSyn pathology segmentation is shown beneath each region, with the $log_{10}$ % area occupied indicated for each region in the representative image Scale bars = 400 μm.

A

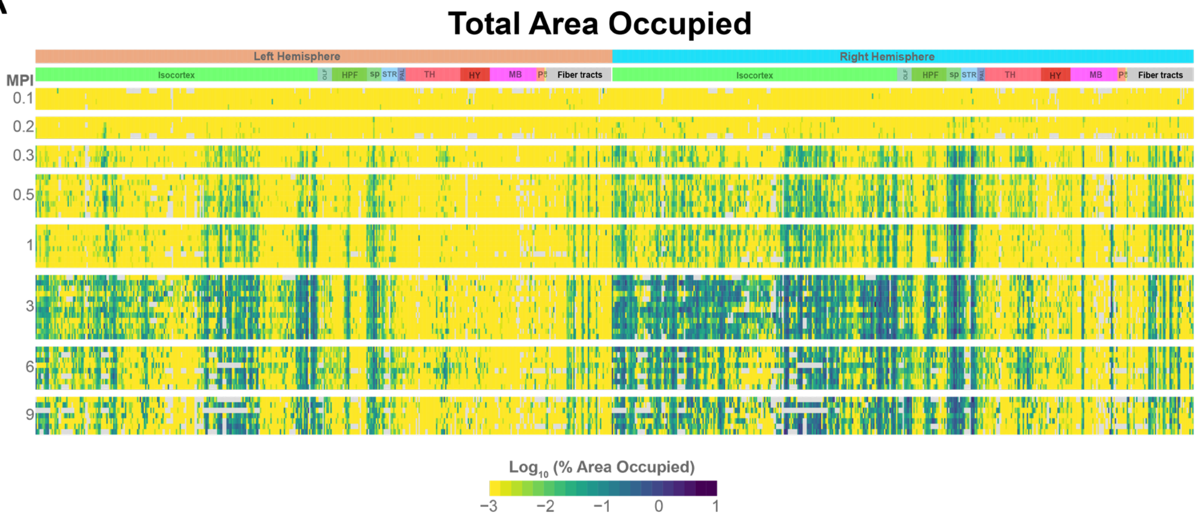
Total Area Occupied
Left Hemisphere
Right Hemisphere
Isocortex
HPF
sp
STR
TH
HY
MB
Fiber tracts
MPI
0.1
0.2
0.3
0.5
1
3
6
9
Log10 (% Area Occupied)
−3 −2 −1 0 1

B

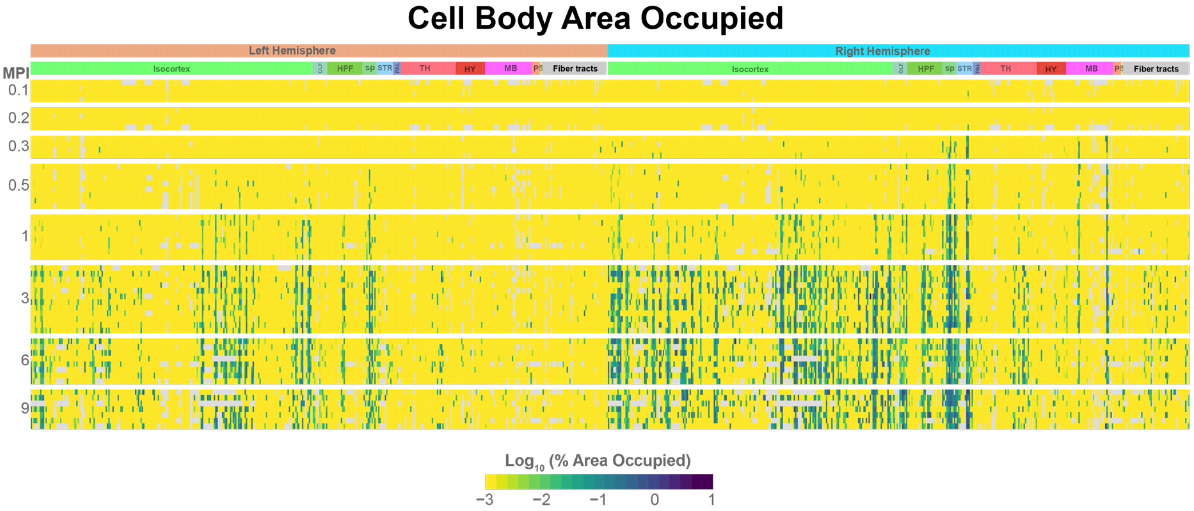
Cell Body Area Occupied
Left Hemisphere
Right Hemisphere
Isocortex
HPF
sp
STR
TH
HY
MB
Fiber tracts
MPI
0.1
0.2
0.3
0.5
1
3
6
9
Log10 (% Area Occupied)
−3 −2 −1 0 1

C

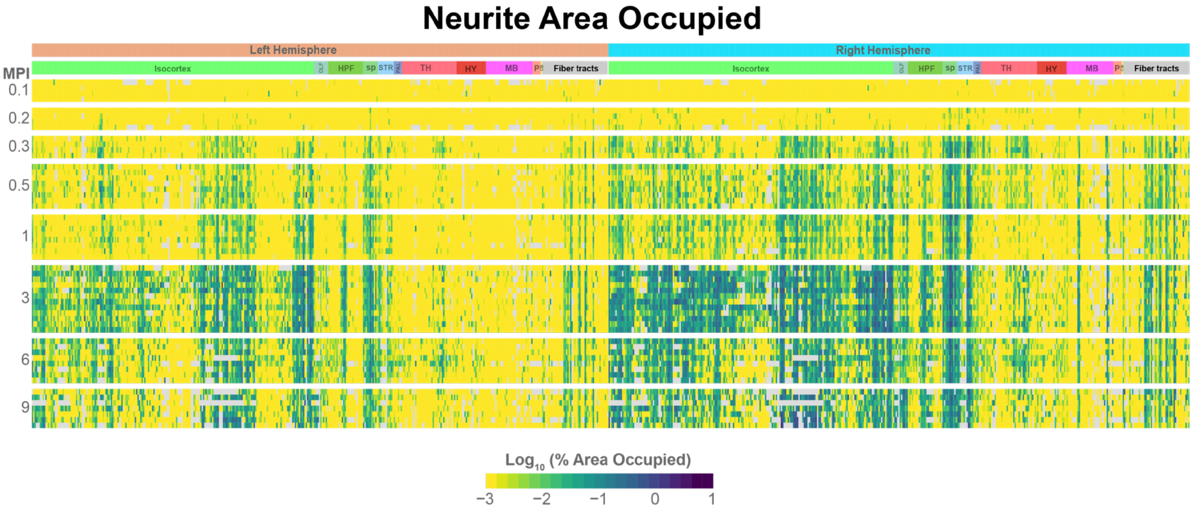
Neurite Area Occupied
Left Hemisphere
Right Hemisphere
Isocortex
HPF
sp
STR
TH
HY
MB
Fiber tracts
MPI
0.1
0.2
0.3
0.5
1
3
6
9
Log10 (% Area Occupied)
−3 −2 −1 0 1

**Figure S3. Grid Heatmaps of total, cell body, and neuritic pathology** (A) Heatmap plot of total regional pathology measures from 0.1-9 MPI. Each row represents an individual mouse, and each column represents a brain region with major regional designations labeled at the top. (B) Heatmap plot of cell body regional pathology measures from 0.1-9 MPI. Each row represents an individual mouse, and each column represents a brain region with major regional designations labeled at the top. (C) Heatmap plot of neuritic regional pathology measures from 0.1-9 MPI. Each row represents an individual mouse, and each column represents a brain region with major regional designations labeled at the top.

### Figure S4

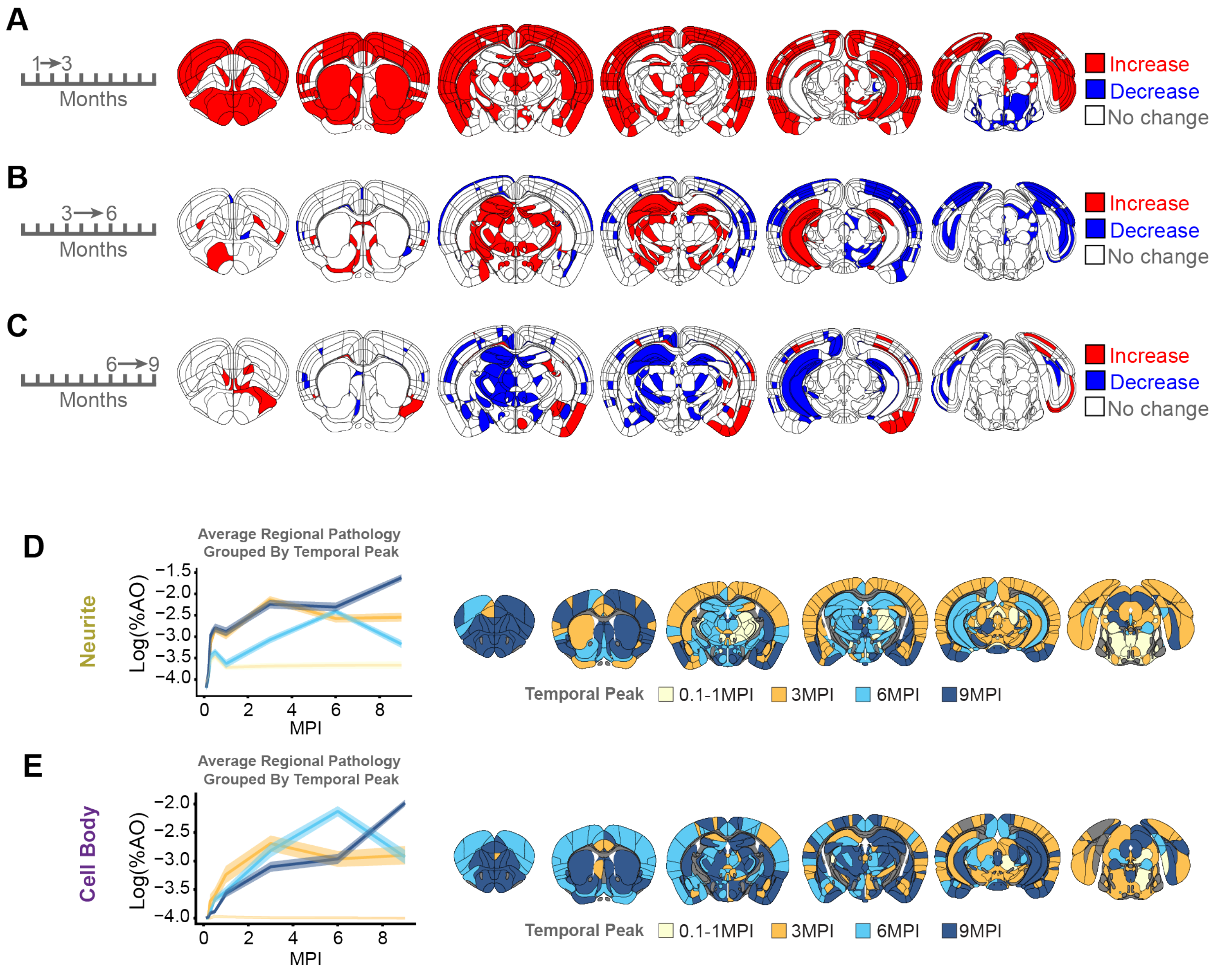


**Figure S4. Time-dependent changes in regional α-synuclein pathology** (A) Statistical comparisons were made comparing α-synuclein pathology in each region from 1 MPI to 3 MPI. Red indicates a statistically-significant increase in pathology (second generation *p* values), while blue indicates a statistically-significant decrease. Most regions have increased pathology at 3 MPI compared to 1 MPI. (B) Comparisons of α-synuclein pathology changes from 3 to 6 MPI. There are many fewer changes, with increases in contralateral hippocampus, but decreases in caudal cortical regions. (C) Comparisons of α-synuclein pathology from 6 to 9 MPI. Almost all regions have peaked and are stable or decreasing, with the exception of some amygdala regions and piriform, which are still increased. (D) Neuritic α-synuclein pathology peaks at different times in specific regional groups. The average pathology level of regions that peak at different times is plotted as a group in the panel on the left. For visualization purposes, regions containing 0 pathology were represented as –4 on the log scale. The anatomical plots on the right display the peak time for each region. Similar data for cell body (E) α-synuclein pathology are also plotted.

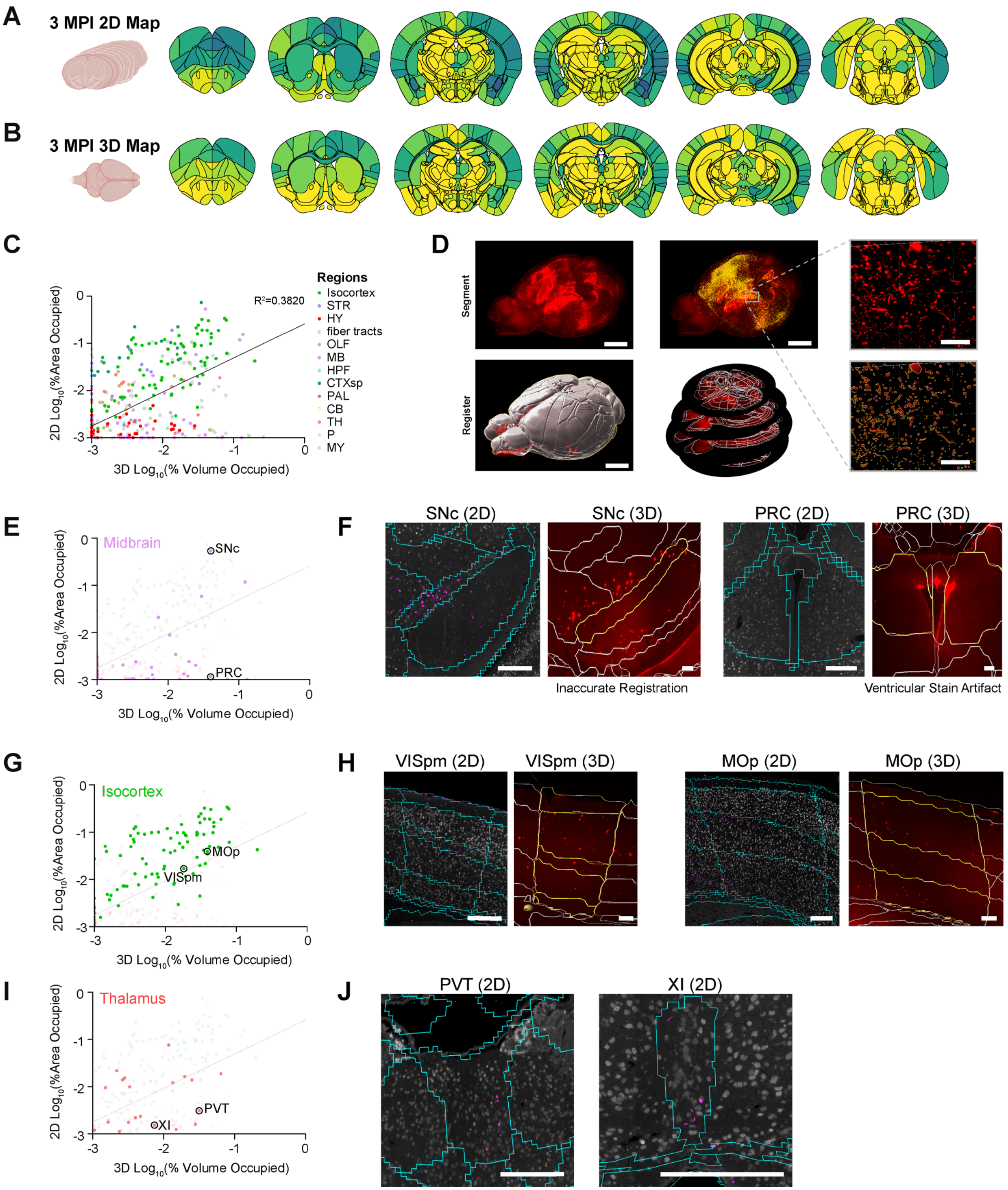


**Figure S5. Comparison of 2D and 3D maps of α-synuclein pathology** (A) Anatomical heatmap representing regional pathology area occupied for 3 MPI mice in our 2D analysis using the modified QUINT workflow. For the comparison to the 3D map, only cortical parent regions (and not laminar data) are shown. (B) Anatomical heatmap representing pathology volume occupied for 3 MPI mice in our 3D analysis. (C) Correlation plot showing individual region pathology in the 2D and 3D analysis. Individual region data points are colored by major regional designations. Linear regression line of best fit and regression co-efficient are plotted. (D) Whole brain analysis methods. 3D reconstruction of mouse brain was rendered in IMARIS. Threshold generated pathology segmentation volume. Registration to the ABA CCFv3 via BrainGlobe registration. Scale bars = 2.5 mm for whole brain volume, 250 μm for zoomed segmentation. (E) Correlation plot highlighting midbrain regions. (F) Selected regions show different pathology levels. In the SNc, the 3D registration led to SNc pathology being mis-assigned to other midbrain regions. In the PRC, no pathology is present, but a ventricular stain artifact leads to aberrant pathology calling in the 3D map. Scale bars = 250 μm. (G) Correlation plot highlighting regions within the isocortex. Cortical regions are generally well-correlated, with generally higher values in the 2D segmentation. (H) Selected visual (VISpm) and motor (MOp) cortical regions show similar amounts of pathology in both 2D and 3D analyses. Scale bars = 250 μm. (I) Correlation plot highlighting thalamic regions. While pathology was generally well-represented in both 2D and 3D analysis, we identified PVT and XI thalamus regions that were not well-sampled by our 2D sampling strategy but were picked up by the 3D map. (J) The pathology in these regions (PVT, XI) was apparent with additional 2D sampling. Scale bars = 250 μm.

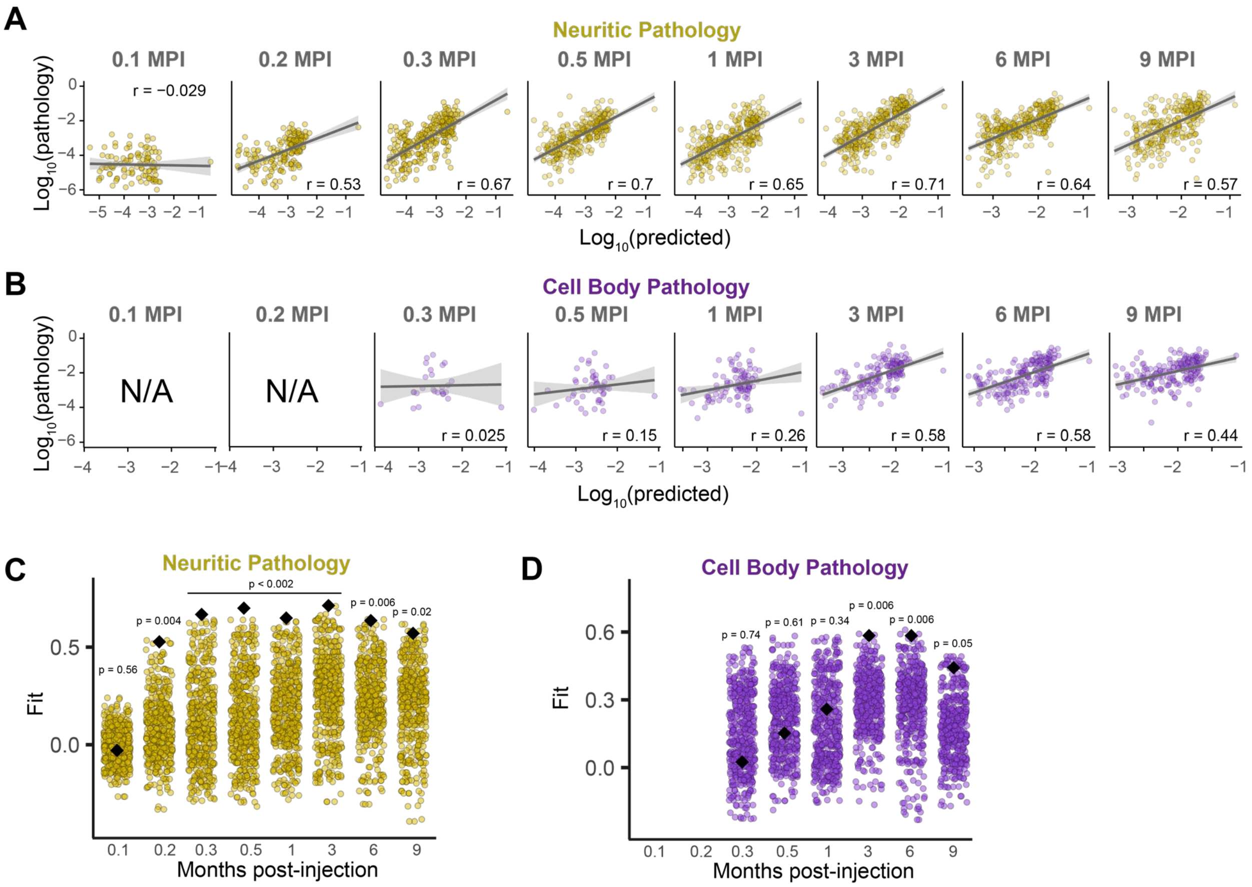


**Figure S6. Computational models of neuritic and cell body pathology based on anatomical connectivity show predictivity of regional α-synuclein pathology** (A) Predictions of neuritic α-synuclein pathology from linear diffusion models based on bidirectional (anterograde and retrograde) anatomical connections. Solid lines represent the line of best fit, and shading represents 95% confidence intervals. (B) Predictions of cell body α-synuclein pathology from linear diffusion models based on bidirectional (anterograde and retrograde) anatomical connections. Solid lines represent the line of best fit, and shading represents 95% confidence intervals. (C) Comparison of Pearson's *r* values obtained by fitting bidirectional spread models using actual (black diamond) and alternate (circles) seed regions for neuritic pathology. (D) Comparison of Pearson's *r* values obtained by fitting bidirectional spread models using actual (black diamond) and alternate (circles) seed regions for cell body pathology.

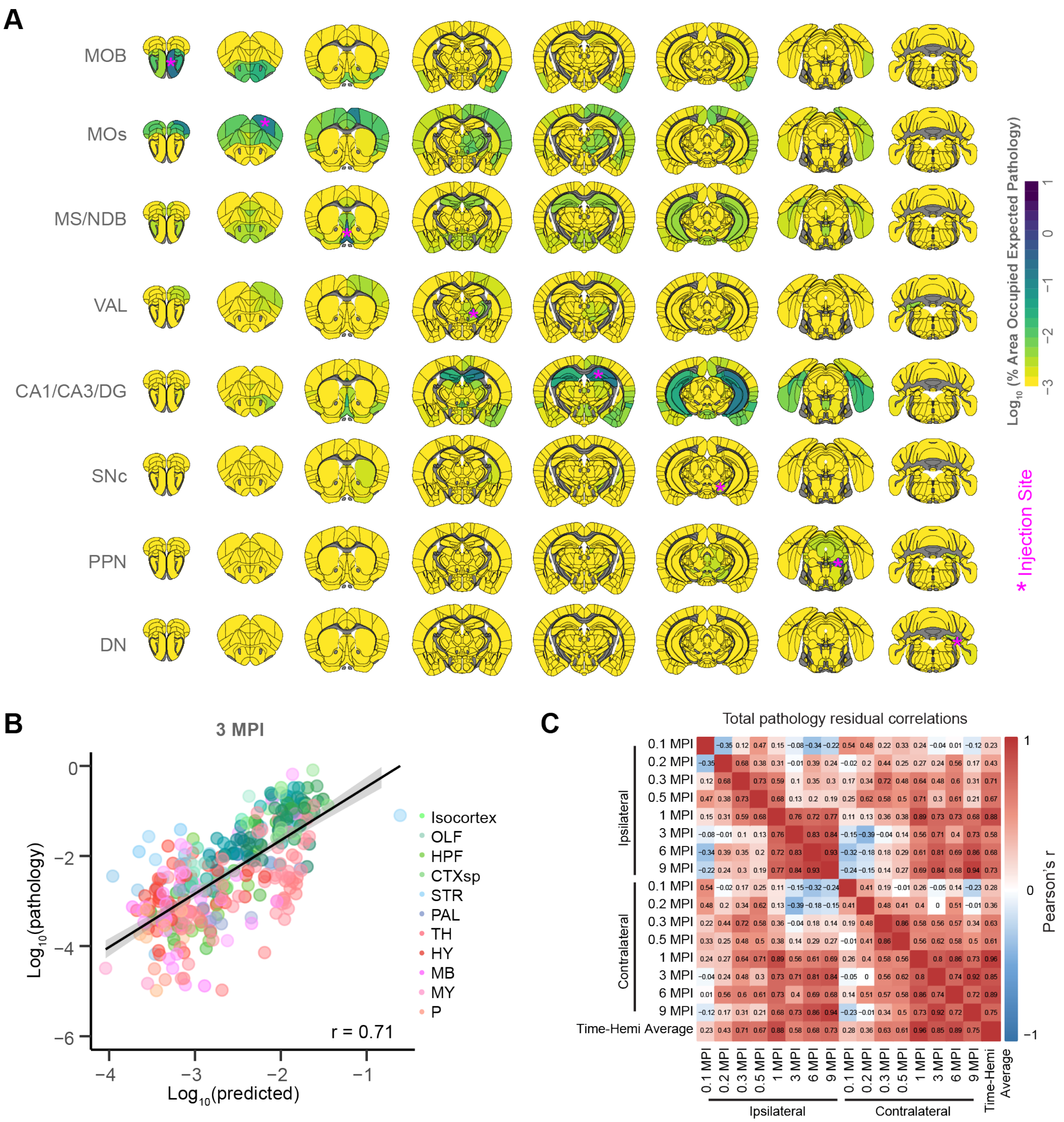


**Figure S7. Computational model predictions of regional vulnerability** (A) Pathology predicted by linear diffusion modeling with the same parameters fit from the caudoputamen injection site, but with alternate injection sites. MOB: main olfactory bulb, MOs: secondary motor area, MS/NDB: medial septal nucleus/diagonal band nucleus, VAL: ventral anterior-lateral complex of the thalamus, CA1/CA3/DG: hippocampus (CA1, CA3, dentate gyrus), SNc: substantia nigra-compact part, PPN: pedunculopontine nucleus, DN: dentate nucleus. (B) Model predictions of total pathology, as in Fig. 4A, plotted with points colored by major anatomical division. This demonstrates that certain anatomical divisions (thalamus, midbrain) have lower pathology than expected based on connectivity. (C) Heatmap displaying the Pearson's correlations between residuals for brain regions derived from computational modeling of α-synuclein pathology at different timepoints and hemispheres. Residuals from 1-9 MPI show strong correlations, and hemispheres are also correlated at these time points.

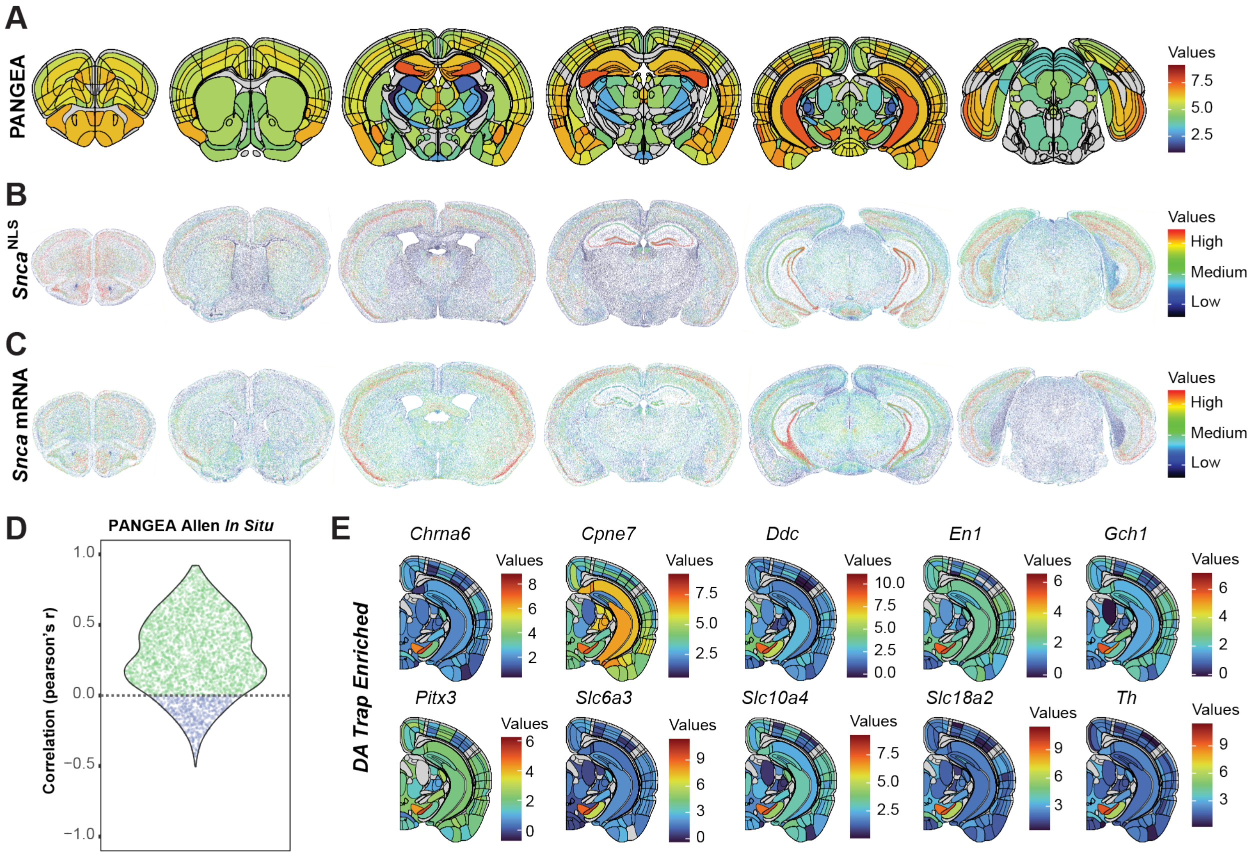


**Figure S8. PANGEA comparisons to other gene expression atlases** (A) Anatomical heatmap of *Snca* expression values in PANGEA compared to (B) protein staining of α-synuclein in mice where α-synuclein is tagged with a nuclear localization signal and (C) *Snca in situ* hybridization staining. Panels B and C display the relative expression of α-synuclein or *Snca* in each cell with warm values corresponding to high expression. Data adapted from PMID: 38504090. (D) Correlation of each gene contained in both Allen *in situ* atlas and PANGEA (Pearson's r). (E) PANGEA anatomical heatmaps of 10 genes previously identified as dopaminergic neuron selective using dopamine-driven RiboTRAP38 or enriched in the substantia nigra over the ventral tegmental area.

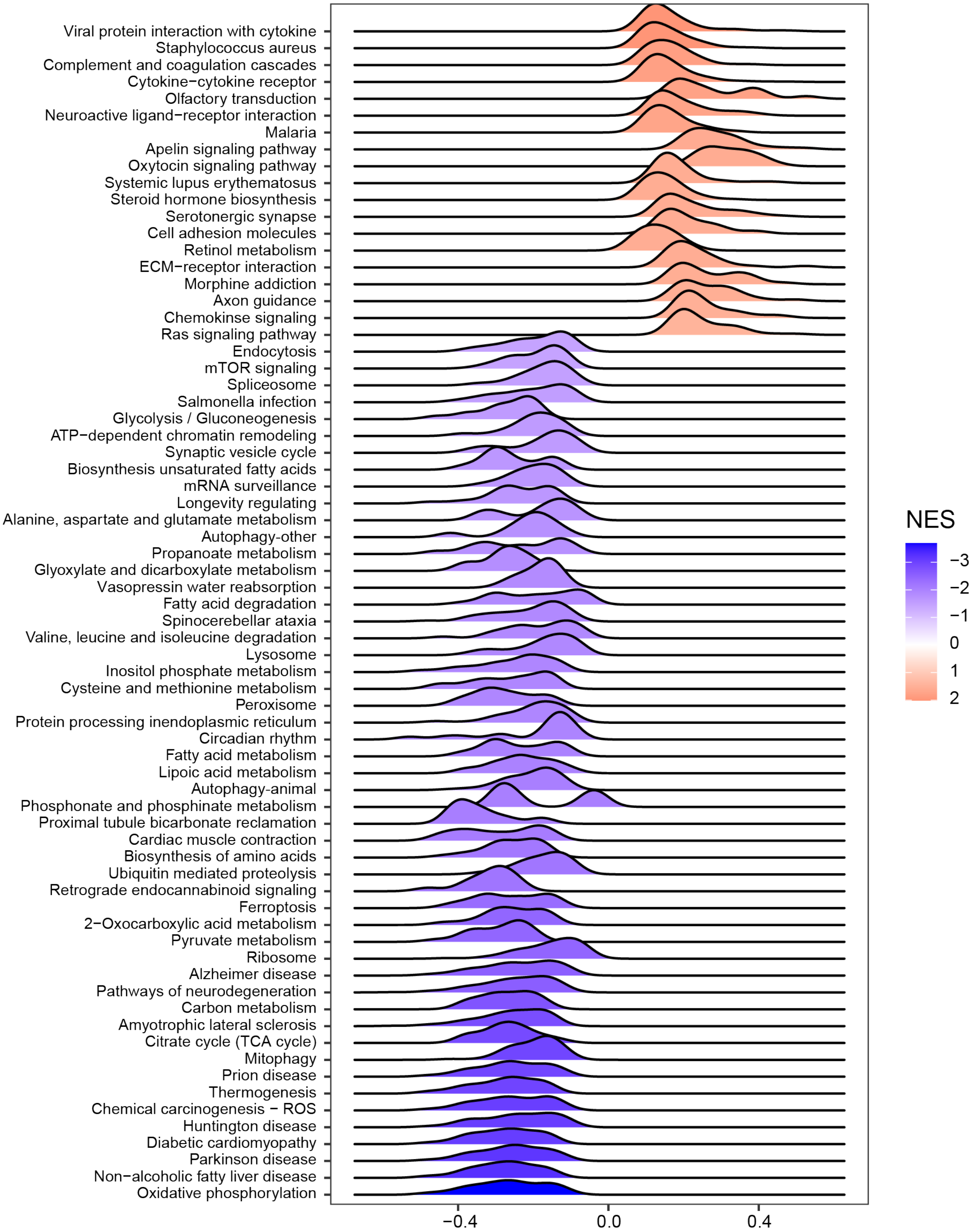


**Figure S9. Gene set enrichment analysis significantly related to regional vulnerability**

Ridgeplots of Normalized Enrichment Scores from gene set enrichment analysis of genes significantly correlated to pathology vulnerability (FDR<0.05)

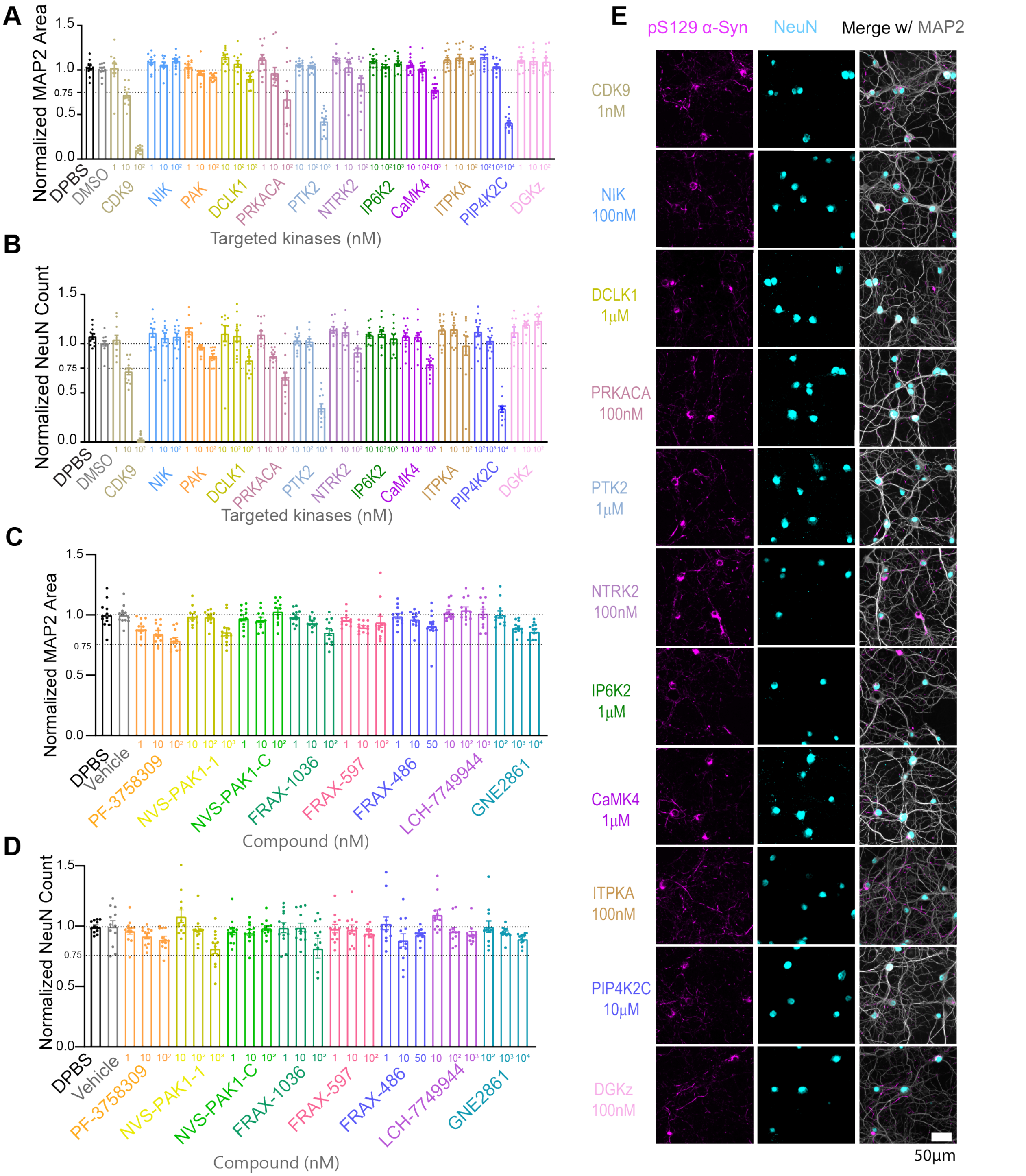


**Figure S10. Effect of kinase inhibitors on neuronal health** (A) MAP2 area and (B) NeuN count normalized to vehicle treated group from the kinases inhibited at 3 different doses in primary hippocampal neurons treated with α-synuclein PFFs from the screen of 12 kinase inhibitors. (C) MAP2 area and (D) NeuN count normalized to vehicle treated group from screen of various PAK inhibitors with varying specificity. Data presented as mean ± SEM with individual values plotted. N=12 independent wells from 4 separate cultures. (E) Representative images of pS129 α-synuclein from the kinase inhibitor screen at the highest safe test dose for each kinase inhibitor. Scale bars = 50 μm.

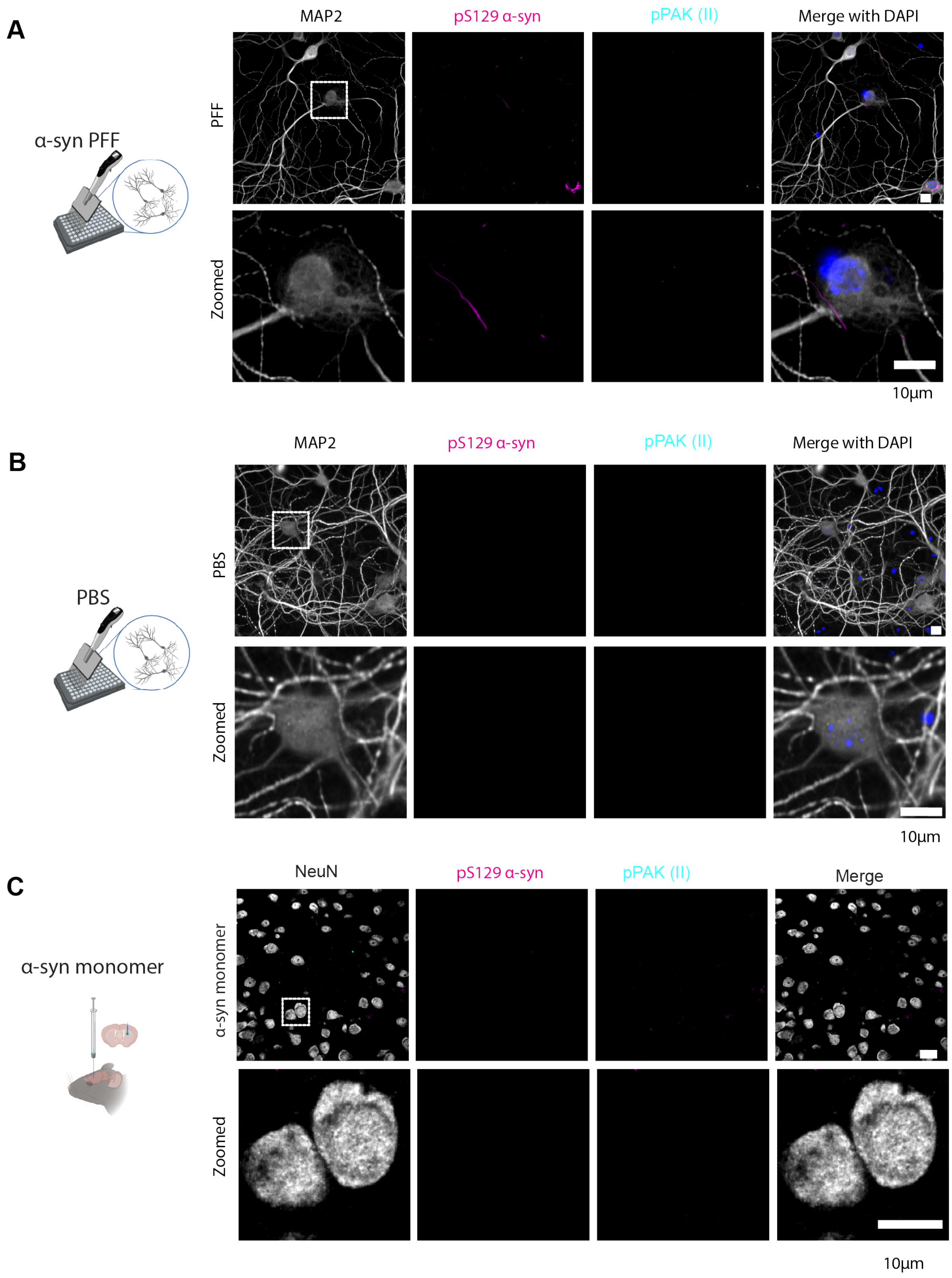


**Figure S11. Group II pPAK in neurons without α-synuclein inclusions**

(A) Representative images of co-immunofluorescence from primary neurons treated with α-synuclein PFFs with a zoom of a neuron without an α-synuclein inclusion. Note, this is the same field of view as Fig. 8H with a zoomed image of a different neuron. (B) Representative images of co-immunofluorescence from primary neurons treated with PBS showing no pPAK (group II) punctate pattern in cells without α-synuclein inclusions. (C) Representative images for co-immunofluorescence of α-synuclein inclusions and pPAK (group II) in amygdala from mouse injected with α-Syn monomer in dorsal striatum and stained 3MPI. Scale bars = 10 μm.

### Figure S12

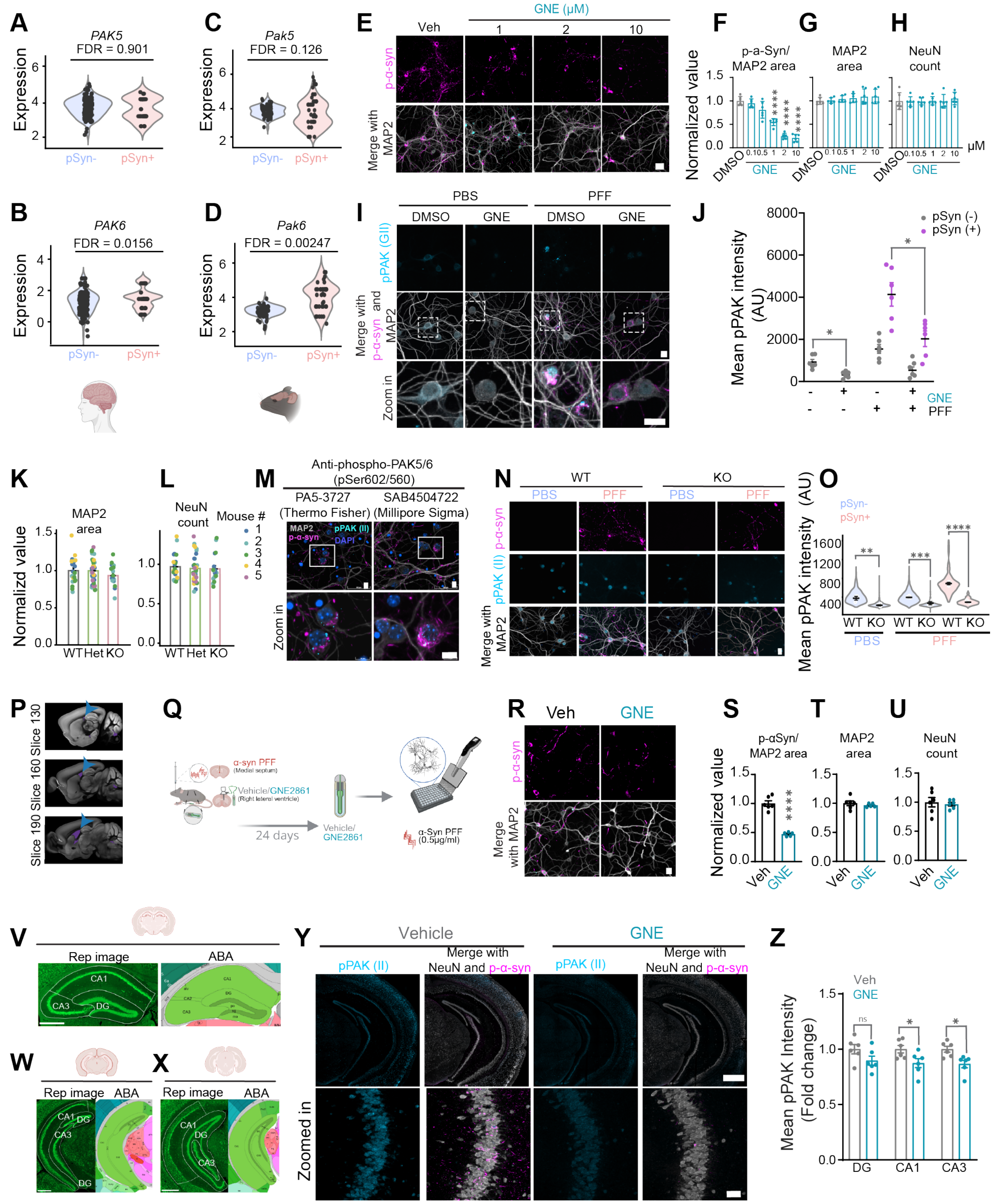

**Figure S12. Supporting validation and experimental rationale for PAK inhibition studies**
(A,B) Quantification of *PAK5* and *PAK6* RNA in α-synuclein inclusion-bearing neurons and non-inclusion-bearing neurons within the same PD brain and in (C,D) α-synuclein PFF injected mice. (E) Representative co-immunofluorescence images from primary neurons treated with α-

synuclein PFFs under low pathology conditions (0.125 μg/mL) and exposed to a range of lower doses of GNE2861. Scale bar = 20 μm. (F) pS129 α-synuclein/MAP2 area (G) MAP2 area and (H) NeuN count normalized to vehicle treated group. Data are presented as mean ± SEM, N=9 independent wells from 3 separate cultures, p-values represent fold-change compared to vehicle control with Welch ANOVA test and Dunnett's T3 multiple comparison test: ****$p < 0.0001$. (I) Representative images and (J) quantification of phospho-PAK (pPAK) intensity in primary neurons treated with preformed fibrils (PFF) and/or GNE2861. Scale bar = 20 μm. Statistical significance was calculated using an unpaired two-tailed t-test with Welch's correction, *$p < 0.05$. Quantification of (K) MAP2 area and (L) NeuN counts normalized to WT controls in primary neurons from Pak5/6 wild type (WT), heterozygous (Het), and knockout (KO) genotypes following PFF treatment. Data are presented as mean ± SEM. (M) Validation of pPAK localization using an independent antibody. Representative images show punctate phospho-PAK (pPAK) signal associated with α-synuclein inclusions. Scale bar =10 μm (N) Representative images of Group II pPAK (PAK5/6) in primary neurons derived from WT and PAK5/6 KO mice. Scale bar = 10 μm. (O) Quantification of pPAK intensity in primary neurons from WT and PAK5/6 KO mice treated with PBS or PFF. Violin plots represent the distribution of pPAK intensity across individual neurons, while individual data points represent individual mice; black bars indicate mean ± SEM. Statistical analysis was performed using two-way ANOVA with genotype and condition as factors, followed by Šídák's multiple-comparisons test comparing WT and KO within each condition. N = 3 per genotype/condition. Adjusted p-values: **$p = 0.0012$ (PBS pSyn−), ***$p = 0.0003$ (PFF pSyn−), and ****$p < 0.0001$ (PFF pSyn+). (P) Representative sagittal mouse brain section from the ABA highlighting the close anatomical proximity of the hippocampus (blue arrows) to the lateral ventricles (magenta) (Q) Experimental schematic for validating efficacy of GNE2861 delivered via ALZET osmotic pumps and recovered at the end of the in vivo study. (R) Representaive immunofluorescence images from primary neurons treated with α-synuclein PFFs and treated with vehicle or GNE2861 (10 μM) from ALZET pumps. Scale bar =20 μm. Quantification of (S) α-synuclein pathology, (T) MAP2 area and (U) NeuN counts normalized to vehicle control. Representative hippocampal sections from experimental animals stained with NeuN corresponding to ABA reference sections (V) Fig. 72, (W) Fig. 82, and (X) Fig. 88. Corresponding ABA maps (atlas.brain-map.org/) indicate the regions of interest (ROIs) used for quantification of pS129 α-synuclein pathology in vehicle- and GNE2861 treated animals following intracerebroventricular (ICV) administration through ALZET osmotic pumps. (Y) Representative hippocampal sections and (Z) quantitation of pPAK (group II) for vehicle- and GNE2861-treated mice injected with α-synuclein PFFs injected corresponding to ABA reference sections Fig 82 . Scale bars = 100 μm. Data represents fold change in Group II pPAK intensity relative to the vehicle control. N = 6 mice per group. Statistical significance was calculated using an unpaired two-tailed t-test with Welch's correction, *$p < 0.05$.
